# Somatic mouse models of gastric cancer reveal genotype-specific features of metastatic disease

**DOI:** 10.1101/2022.06.15.494941

**Authors:** Josef Leibold, Corina Amor, Kaloyan M. Tsanov, Yu-Jui Ho, Francisco J. Sánchez-Rivera, Judith Feucht, Timour Baslan, Hsuan-An Chen, Sha Tian, Janelle Simon, Alexandra Wuest, John E. Wilkinson, Scott W. Lowe

## Abstract

Metastatic gastric carcinoma is a highly lethal cancer that responds poorly to conventional and molecularly targeted therapies. Despite its clinical relevance, the mechanisms underlying the behavior and therapeutic response of this disease are poorly understood owing, in part, to a paucity of tractable models that faithfully recapitulate different subtypes of the human disease. To close this gap, we developed methods to somatically introduce different oncogenic lesions directly into the stomach epithelium and show that genotypic configurations observed in patients produce metastatic gastric cancers that recapitulate the histological, molecular, and clinical features of all non-viral molecular subtypes of the human disease. Applying this platform to both wild-type and immune-deficient mice revealed previously unappreciated links between the genotype, organotropism and immune surveillance of metastatic cells that produced distinct patterns of metastasis that were mirrored in patients. Our results establish and credential a highly portable platform for producing autochthonous cancer models with flexible genotypes and host backgrounds, which can unravel mechanisms of gastric tumorigenesis or test new therapeutic concepts aimed at improving outcomes in gastric cancer patients.

## INTRODUCTION

Gastric cancer is the fourth leading cause of cancer-associated deaths and the fifth most commonly diagnosed cancer worldwide^1^. While localized disease can be successfully treated, the survival rate of gastric cancer patients drops dramatically in the advanced and especially at the metastatic stages^2–4^. Despite recent advances in our understanding of the molecular features of this cancer, effective treatment strategies are currently lacking, particularly for patients with metastatic disease^5, 6^.

Genome sequencing studies have classified gastric cancer into four major molecular subtypes defined by: (1) chromosomal instability (CIN); (2) genomic stability (GS); (3) microsatellite instability (MSI); and (4) Epstein-Barr Virus (EBV) infection^7–9^. The CIN tumor subtype is the largest and typically harbors *TP53* mutations and a high frequency of recurrent copy number alterations (CNAs)^10^. In contrast, tumors of the GS subtype display far fewer chromosomal aberrations and are devoid of *TP53* mutations, instead frequently harboring mutations that inactivate *CDH1* or activate the WNT signaling pathway. Not only do CIN and GS tumors display distinct mutational profiles, but they also differ in their histopathology, showing prominent features of intestinal differentiation or diffuse histological features, respectively^11^. The MSI subtype is defined by the presence of microsatellite instability and mutations in mismatch repair genes such as *MLH1* or *MSH2*. Presumably due to their increased mutational load and potential for neoantigen production, these tumors elicit a T cell-dominated immune response^12, 13^ and frequently respond to immune checkpoint blockade^12, 14–17^. Finally, the EBV tumor subtype is a uniquely different entity that is characterized by alterations in the PI3K/AKT signaling pathway^10, 18^ and typically displays a diffuse and intestinal histopathology^19^. Interestingly, mutational gains and amplifications of the *MYC* gene are associated with early progression of intestinal metaplasia to gastric cancer and can be found in all gastric cancer subtypes^7, 10^.

Owing to their ability to capture tumor development in the complexity of the whole organism, genetically engineered mouse models (GEMMs) have proven to be a valuable tool for understanding genotype-phenotype relationships and evaluating new therapeutic concepts in a range of tumor types. However, due to the cost and waste of intercrossing various germline strains, traditional GEMMs are time and resource-consuming, making it difficult to model and interrogate the spectrum of tumor genotypes that exists in patients or conduct large-scale preclinical studies^20–23^. Likewise, it is extremely cumbersome to interrogate the genetics of tumor-host interactions, as the number of intercrosses needed to produce a genetically defined cancer and in an altered host strain is prohibitive. For gastric cancer, existing GEMMs only model some molecular subtypes on a single host background and, in contrast to patients, rarely progress to metastatic disease^24^. Therefore, the availability of new models that capture the genetic diversity and metastatic progression of human gastric cancer and enable facile changes in the host would transform the study of this disease.

We and others have devised methods to somatically introduce cancer predisposing lesions or other genetic elements into murine tissues using electroporation, thereby producing <u>E</u>lectro<u>po</u>ration-based <u>G</u>enetically <u>E</u>ngineered <u>M</u>ouse <u>M</u>odels (EPO-GEMMs)^25,26,27,28^. In this approach, transposon-based vectors encoding cDNAs or CRISPR-Cas9 constructs targeting endogenous genes are introduced into the tissue via survival surgery through a brief electric pulse, whereby they are taken up by a subset of cells. In circumstances where a particular lesion or combination of lesions provides a selective advantage, focal tumors arise at the electroporation site. Herein, we developed surgical methods and electroporation conditions suitable for engineering mice with gastric tumors harboring a range of cancer genotypes and show that the resulting platform can faithfully model the three major non-viral subtypes of the human disease. Furthermore, we illustrate the power of combining this approach with mice of different genetic backgrounds to explore tumor-host interactions relevant to metastatic spread. The portability, flexibility, and speed of these gastric EPO-GEMMs creates new possibilities for exploring how gastric cancers evolve, spread, and respond to therapy in the context of the complex *in vivo* environment.

## RESULTS

### Modeling CIN and GS subtypes of gastric cancer through somatic tissue engineering

To generate gastric cancer EPO-GEMMs, we developed a survival surgery technique coupled with direct tissue electroporation to deliver genetic elements to the murine stomach epithelium (see Methods). A transposase-transposon vector pair was used to express a defined oncogene and a plasmid co-expressing Cas9 with an sgRNA to knock out a tumor suppressor gene of interest (**Fig. 1A**). Since *MYC* is a potent oncogene that is frequently amplified across gastric cancers^7, 10^, we used a transposon vector containing human *MYC* cDNA as the universal oncogene, and adapted sgRNAs to target different tumor suppressor genes in accordance with their mutation in distinct subtypes of gastric cancer (**Fig. 1B**).

**Figure 1.**
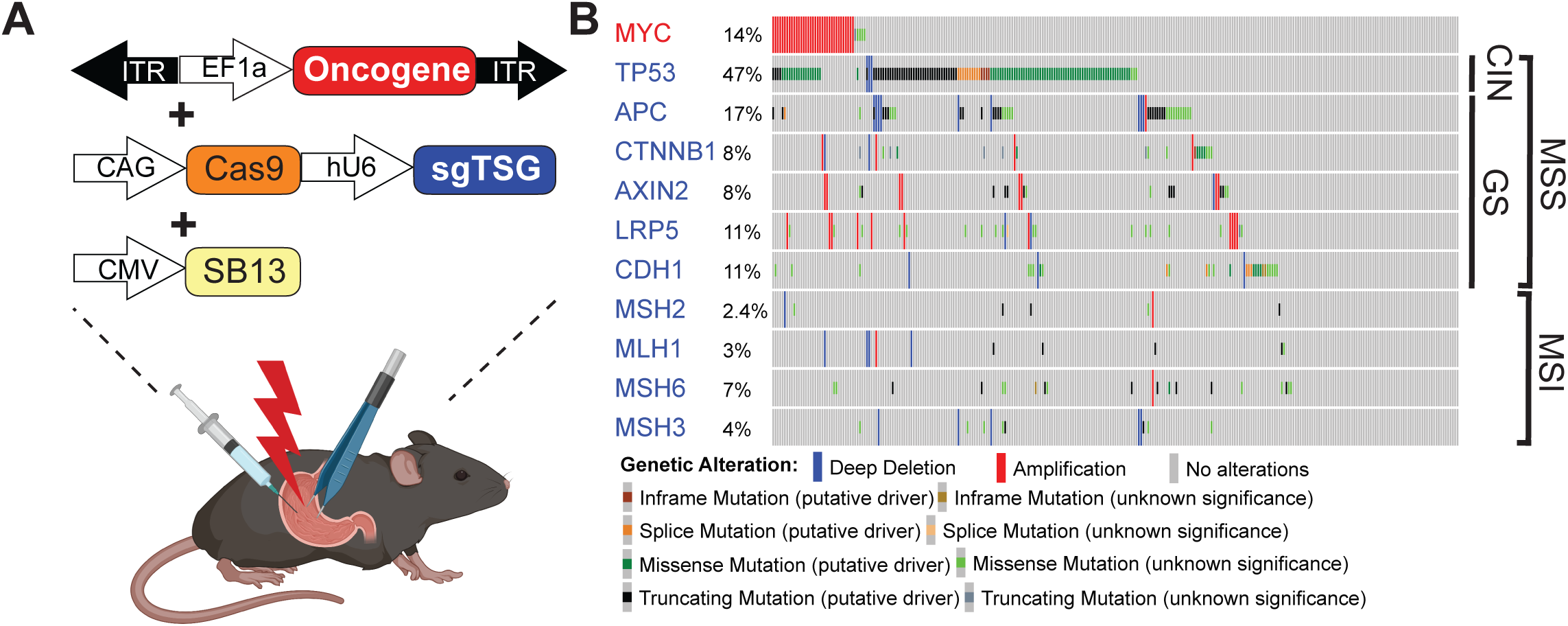
Modeling molecular subtypes of gastric cancer in mice by a somatic tissue engineering approach. (A) Schematic of the <u>e</u>lectro<u>po</u>ration-induced <u>g</u>enetically <u>e</u>ngineered <u>m</u>ouse <u>m</u>odels (EPO-GEMMs) of gastric cancer. A transposon vector harboring an oncogene in combination with a Sleeping Beauty transposase (SB13) and/or a CRISPR-Cas9 vector targeting tumor suppressor genes (TSGs) are delivered into the stomach by direct *in vivo* electroporation. (B) MSK-IMPACT oncoprint displaying the genomic status of recurrent oncogenes and tumor suppressor genes in gastric cancer patients. Associated molecular subtypes (per TCGA^10^) are shown on the right.

We first set out to develop a model of the most common subtype observed in patients, CIN gastric cancer, which is characterized by a high frequency of *TP53* mutations^8, 10^. The *MYC* transposon-transposase system was combined with a Cas9-sgRNA vector targeting *Trp53* (hereafter referred to as *p53*) to recapitulate a genotype commonly seen in patients^29^ (**Fig. 1B**). Mice electroporated with all three plasmids consistently developed lethal tumors (90% penetrance; 45 days medium survival) that harbored the predicted disruptions of the *p53* locus (**Fig. 2A; Extended Data Fig. 1A**). In contrast, mice electroporated with either the *MYC* or Cas9-sgp53 vector alone did not develop tumors within one year of follow-up (**Fig. 2A**). The resulting tumors displayed a mixed histopathology, consisting predominantly of well-differentiated adenocarcinoma of the intestinal phenotype^11^ and regions of poorly differentiated gastric carcinoma that were present in late-stage tumors (**Fig. 2B**). The well-differentiated areas expressed E-cadherin, cytokeratin-8 (CK8), high levels of the proliferation marker Ki67, and partially stained positive for the parietal cell marker H^+^/K^+^ ATPase, in accordance with human CIN gastric tumors (**Fig. 2B; Extended Data Fig. 1B**). These observations are consistent with an epithelial cell of origin of the EPO-GEMM tumors, which was confirmed by generating tumors with comparable latency and presentation in a CK8-CreERT2; LSL-Cas9 host that restricts tumor initiation to the CK8^+^ epithelial compartment (**Extended Data Fig. 2A-C**).

**Figure 2.**
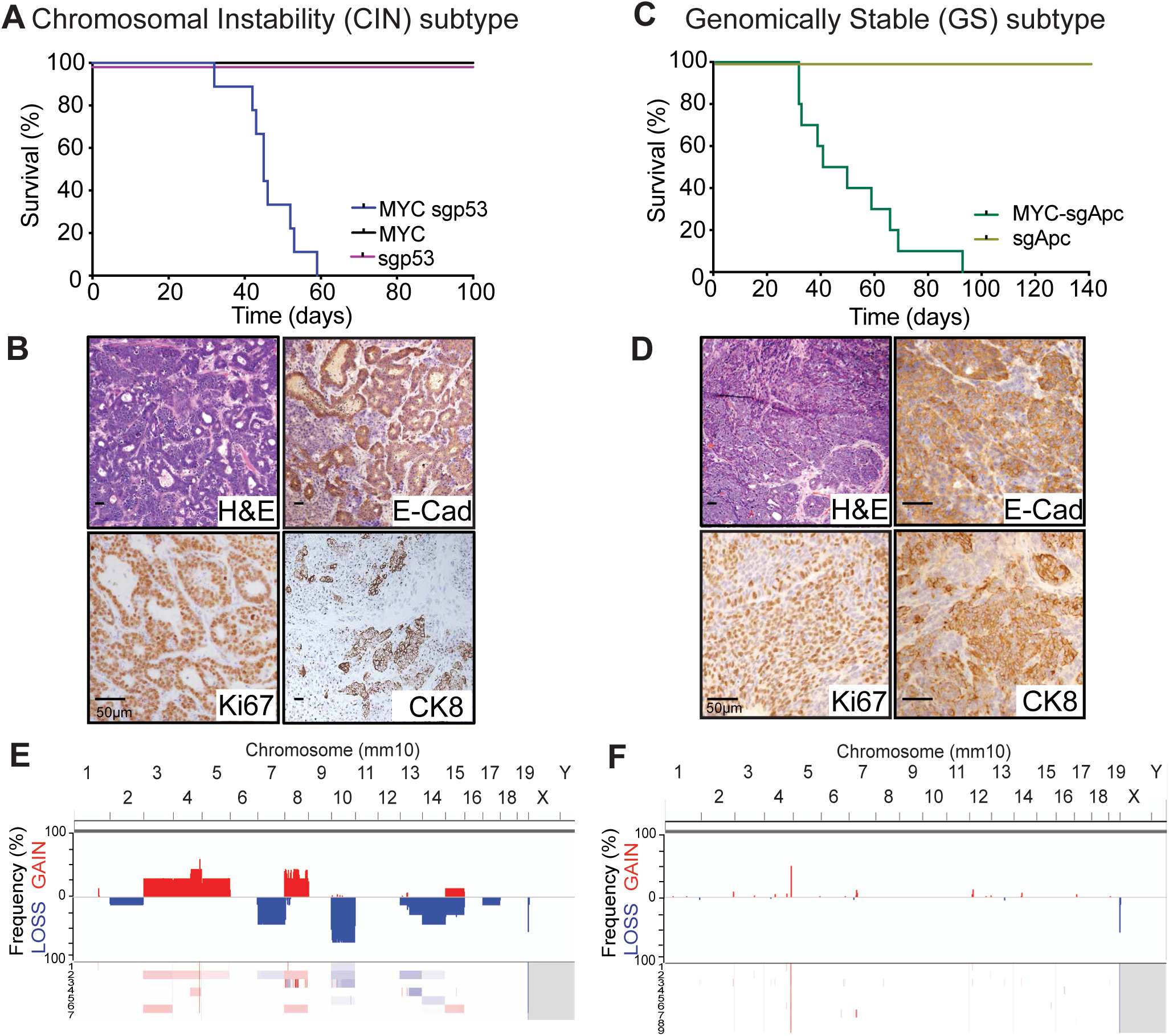
CIN and GS gastric cancer EPO-GEMMs recapitulate hallmark histological and molecular features of the corresponding human subtypes. (A) Kaplan-Meier survival curve of C57BL/6N mice electroporated with: a *MYC* transposon vector and a Sleeping Beauty transposase (*MYC*; black, n=4); a CRISPR-Cas9 vector targeting *p53* (sgp53; red, n=4); or the combination of all vectors (*MYC* sgp53; blue, n=9). (B) Representative hematoxylin and eosin (H&E) and immunohistochemical staining for E-cadherin (E-Cad), Ki67 and cytokeratin 8 (CK8) of a *MYC-p53^-/-^* gastric EPO-GEMM tumor. (C) Kaplan-Meier survival curve of C57BL/6 mice electroporated with: a CRISPR-Cas9 vector targeting *Apc* (sgApc; grey, n=3); the combination of Sleeping Beauty, MYC and a CRISPR-Cas9 vector targeting *Apc* (*MYC* sgApc; green, n=10). (D) Representative hematoxylin and eosin (H&E) and immunohistochemical staining for E-cadherin (E-Cad), Ki67 and cytokeratin 8 (CK8) of a *MYC-Apc^-/-^* gastric EPO-GEMM tumor. (E-F) sWGS analysis of copy number alterations in *MYC-p53^-/-^* (n=7) (E) and *MYC-Apc^-/-^* (n=9) (F) gastric EPO-GEMM tumors. Frequency plots are shown on the top and individual sample tracks are provided on the bottom.

Next, we proceeded to model the GS subtype of gastric cancer. Since human GS tumors frequently harbor alterations in WNT pathway genes and/or *CDH1* (encoding E-cadherin) (**Fig. 1B**), we replaced the *p53* sgRNA with an sgRNA targeting *Apc* or *Cdh1* (**Extended Data Fig. 1C, E**). Delivery of the MYC-sgCdh1 or MYC-sgApc configurations to the gastric epithelium consistently produced tumors with a median survival of 68 and 44 days post-electroporation, respectively (**Fig. 2C; Extended Data Figs. 1F, 2E**). Histological characterization of the *MYC-Apc^-/-^* tumors revealed an undifferentiated histology while largely retaining expression of the epithelial markers E-cadherin and CK8, and partially stained positive for H^+^/K^+^ ATPase, as seen in human gastric cancer (**Fig. 2D; Extended Data Figs. 1D, 2F**). On the other hand, the *MYC-Cdh1^-/-^* tumors displayed undifferentiated histology that, as expected, showed a complete absence of E-cadherin expression (**Extended Data Fig. 1G**). Interestingly, this diffuse undifferentiated histopathology was remarkably similar to that arising in late-stage CIN tumors, which also became E-cadherin negative (**Extended Data Fig. 2C-D**). These observations raise the possibility that p53-associated cell plasticity is an important element of tumor evolution in CIN tumors and, in agreement, we noted that *TP53* and *CDH1* mutations are mutually exclusive in human gastric cancer patients (**Extended Data Fig. 2G)**.

To molecularly characterize the chromosomal stability of CIN and GS EPO-GEMM tumors, we performed sparse whole-genome sequencing of *MYC-p53^-/-^*, *MYC-Apc^-/-^*, and *MYC-Cdh1^-/-^* tumors. Importantly, *MYC-p53^-/-^* but not *MYC-Apc^-/-^*or *MYC-Cdh1^-/-^* tumors harbored recurrent genomic rearrangements that showed synteny to their human counterparts, consistent with the CIN subtype of human gastric cancer (**Fig. 2E-F; Extended Data Fig. 1H-I**). Taken together, the above data establish gastric cancer EPO-GEMMs as fast and flexible models that recreate fundamental histological and molecular features of the CIN and GS subtypes of the human disease.

### Somatic loss of Msh2 induces MSI gastric cancer in mice

Alterations in DNA mismatch repair genes are frequently found in patients and lead to microsatellite instability (MSI) gastric cancer, a subtype that is characterized by an increased frequency of mutations^13, 30^ and a particular base substitution signature^31^ that has not been produced using traditional GEMMs^8, 10^ (**Fig. 1B**). To generate such models, we combined the *MYC* transposon-transposase system with a CRISPR vector co-targeting *p53* and the mismatch repair gene *Msh2*. This approach allows for direct comparison of MSI (*MYC-p53^-/-^-Msh2^-/-^*) and MSS (*MYC-p53^-/-^*) gastric cancers that harbor identical driver genes except for *Msh2* loss.

Consistent with the less aggressive nature of MSI compared to MSS tumors in human gastric cancer patients^8^, the median survival of mice electroporated with *Msh2* sgRNAs was longer compared to *MYC-p53^-/-^* controls (53 vs. 45 days, respectively) (**Fig. 3A**). Importantly, despite this extended survival, *Msh2* disruption appeared to confer a selective advantage during tumorigenesis, as the resulting tumors harbored genetic alterations of the *Msh2* locus and lacked Msh2 expression in the tumor (**Fig. 3B**; **Extended Data Fig. 3A**). These tumors again displayed a mixture of well-differentiated E-cadherin expressing adenocarcinoma and poorly differentiated gastric carcinoma at late-stage disease (**Fig. 3B**). Furthermore, whole-exome sequencing of EPO-GEMM tumors revealed a significantly higher number of genetic alterations in MSI vs. MSS tumors, mainly consisting of single nucleotide variants, small indels (mostly of a single base pair), and a C>T and T>C dominated base substitution signature consistent with human MSI cancers^31^ (**Fig. 3C-D; Extended Data Fig. 3B**).

**Figure 3.**
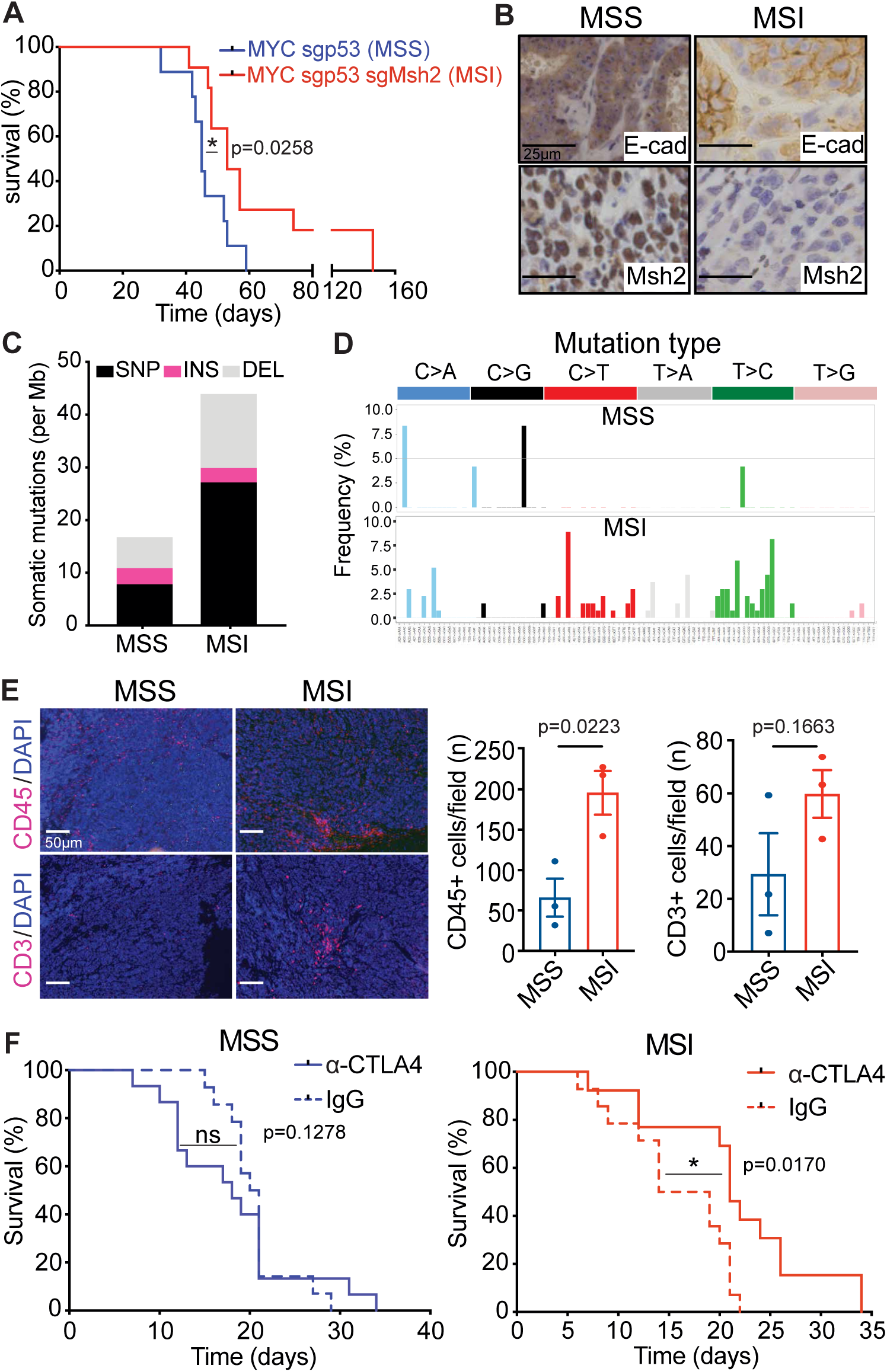
Somatic loss of *Msh2* induces microsatellite instability (MSI) gastric cancer in mice. (A) Kaplan-Meier survival curve of C57BL/6 EPO-GEMMs with either *MYC-p53^-/-^* (MSS; same cohort as shown in Fig. 2A; blue, n=9) or *MYC-p53^-/-^-Msh2^-/-^* (MSI; red, n=11) gastric cancer. (B) Representative immunohistochemical staining for E-cadherin and Msh2 of *MYC-p53^-/-^* (MSS) or *MYC-p53^-/-^-Msh2^-/-^*(MSI) gastric EPO-GEMM tumors. (C) WES analysis of somatic mutations per Megabase (Mb) in either *MYC-p53^-/-^* or *MYC-p53^-/-^-Msh2^-/-^* gastric EPO-GEMM tumors (n=3 independent mice each). SNP = single nucleotide polymorphisms; INS = insertions; DEL = deletions. (D) Base substitution signature in *MYC-p53^-/-^*(MSS) and *MYC-p53^-/-^-Msh2^-/-^* (MSI) gastric EPO-GEMM tumors (n=3 independent mice each). (E) Representative immunofluorescence staining of *MYC-p53^-/-^* (MSS) or *MYC-p53^-/-^-Msh2^-/-^*(MSI) gastric EPO-GEMM tumors for CD45 (red, upper panel) or CD3 (red, lower panel). Quantification to the right (n=3 independent mice each). (F) Kaplan-Meier survival curve of C57BL/6 gastric cancer EPO-GEMMs of either *MYC-p53^-/-^* (left) (n=14 IgG treated, 15 9H10 treated) or *MYC-p53^-/-^-Msh2^-/-^* (right) (n=14 IgG treated, 12 9H10 treated) genotype after antibody-mediated blockade of CTLA-4 (9H10, 200µg) (solid line) or IgG control (dashed line). Treatment was initiated (day 0) after tumor formation was confirmed by abdominal palpation. Statistical analysis: (A), (F) Log-rank test; (E) Unpaired t test.

The high mutational burden of MSI tumors has been shown to result in an increased level of tumor neoantigens presented on MHC class I molecules that can facilitate a T cell-mediated anti-tumor response^13, 32^ and contribute to their increased responsiveness to immune-modulatory drugs^12, 14, 15, 33^. Accordingly, we observed an overall increase of infiltrating CD45^+^ and CD3^+^ cells in MSI EPO-GEMMs compared to their MSS counterparts albeit with substantial intra-tumoral heterogeneity, possibly reflecting the random process of generating immunogenic neoantigens (**Fig. 3E**). Consistent with observations in gastric cancer patients^15^, MSI but not MSS tumors also responded to anti-CTLA4-mediated checkpoint blockade (**Fig. 3F; Extended Data Fig. 3C**). Therefore, these MSI EPO-GEMMs recapitulate the genetic, microenvironmental, and therapeutic response patterns of human MSI gastric cancers.

### EPO-GEMMs recapitulate transcriptional features of human gastric cancer subtypes

Human gastric tumors exhibit gene expression patterns that reflect features of their molecular classification^10^. Hence, we performed bulk RNA sequencing on tumors from EPO-GEMMs that represent the GS, CIN and MSI subtypes, as well as on normal gastric tissue. Hierarchical clustering of all samples indicated that tumor genotype was the most prominent factor dictating the transcriptional landscape of different tumors (**Fig. 4A**). Consistent with human data^10^ and the role of p53 loss in increasing plasticity^34^ **(Fig. 2B; Extended Data Fig. 2)**, CIN tumors showed the greatest inter-tumoral heterogeneity. Gene Ontology analysis of six clusters that segregated differentially expressed genes across all samples revealed transcriptional features that were either tumor-universal or tended to group with specific tumor subtypes **(Fig. 4A; Extended Data Table 1)**. First, as expected for transformed tissue, there was an enrichment of proliferation-related and depletion of differentiation-related pathways across all tumor samples. Second, GS tumors showed a prominent WNT signaling signature, consistent with their *Apc*-null status. Third, CIN tumors exhibited a weak but statistically significant enrichment of extracellular matrix (ECM) genes, which may reflect p53-related ECM remodeling seen in other cancers^35–37^. Fourth, in agreement with our immune-focused analysis above, MSS tumors under-expressed genes involved in inflammatory signaling pathways, as well as genes involved in metabolism and vesicular transport. On the other hand, MSI tumors showed reduced expression of genes involved in oxidative phosphorylation, perhaps due to mitochondrial damage linked to mismatch repair deficiency^38, 39^.

**Figure 4.**
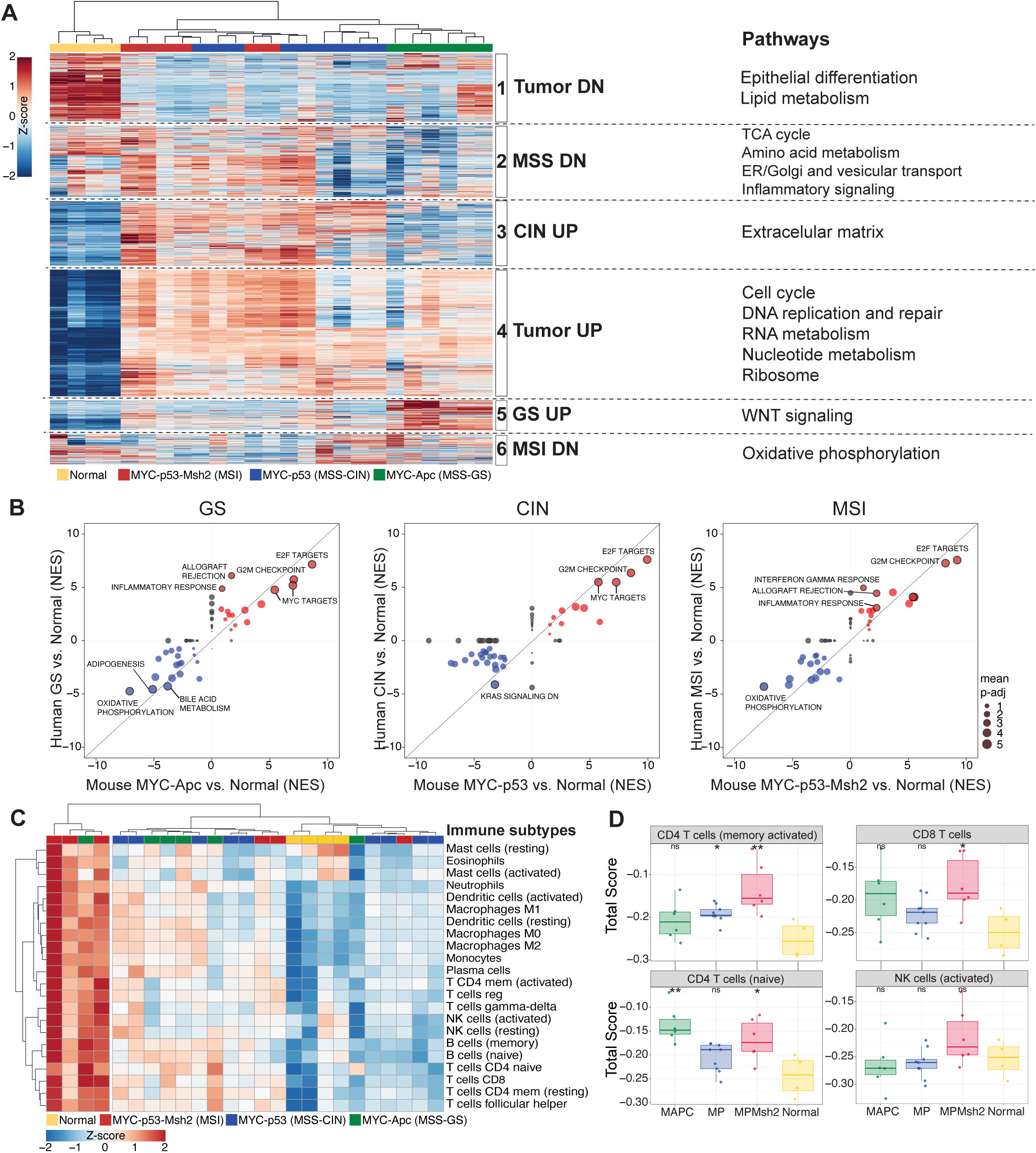
EPO-GEMMs recapitulate transcriptional features of human gastric cancer subtypes. (A) Heatmap of differentially expressed genes across indicated EPO-GEMM samples. Hierarchical clustering segregated all samples based on six signatures (1-6). Key pathways enriched in each signature are shown on the right. Complete lists of genes and pathway predictions are provided in **Extended Data Table 1**. (B) Comparison of GSEA normalized enrichment scores (NES) for Hallmark Pathways enriched in EPO-GEMM (x-axis) and human (y-axis) tumors vs. normal stomach for the indicated genotypes/subtypes. Key pathways are highlighted. Circle size represents the adjusted p-value. Complete lists of pathways and NES scores are provided in **Extended Data Table 1**. (C) Heatmap of CIBERTSORT signatures for distinct immune subpopulations in the indicated EPO-GEMM tumor and normal gastric samples. (D) Box-plots of CIBERSORT signature scores for the indicated immune populations and EPO-GEMM samples. Complete lists of are provided in **Extended Data Table 1**. *p<0.05; **p<0.01, ns=non-significant, Wilcoxon signed-rank test.

These observations were reinforced by Gene Ontology analysis of shared and unique differentially expressed genes for the distinct tumor genotypes, which highlighted a relative depletion of p53 signatures in *MYC-p53^-/-^* tumors and MSI-specific enrichment of immune-related pathways **(Extended Data Fig. 4A, B, F; Extended Data Table 2)**. Importantly, the transcriptional features of EPO-GEMM tumors correlated well with those of human gastric tumors of the respective subtypes, which was largely driven by dominant MYC, proliferation, and immune-related signatures **(Fig. 4B; Extended Data Fig. 4C-E; Extended Data Tables 1, 2).**

We also explored the nature of immune cell infiltrates in different tumor subtypes, which takes advantage of an algorithm known as CIBERSORT^40, 41^ to identify immune cell signatures in bulk tumor samples. Hierarchical clustering segregated a subset of MSI tumors (3/6 samples) as broadly enriched in most immune signatures (**Fig. 4C**), including CD4^+^ and CD8^+^ T cells **(Fig. 4D; Extended Data Table 1)**. Importantly, the immune cell infiltrates and associated signaling pathways displayed marked similarities between murine and human MSS vs. MSI tumors, with the latter showing increased expression of inflammatory pathways and most immune cell signatures (**Extended Data Fig. 4G**). Overall, these gene expression data further demonstrate the molecular fidelity of EPO-GEMMs to their human counterparts and provide insights into pathways that may underlie both common and unique features of different gastric cancer subtypes.

### EPO-GEMMs recapitulate the metastatic organotropism of human gastric cancer

Perhaps the most clinically important feature of gastric cancer is its propensity to metastasize, a property that is rarely observed in traditional gastric cancer GEMMs^22, 24^. In contrast, gastric cancer EPO-GEMMs reproducibly presented with distant metastasis to the liver and lung, as frequently observed in human patients (**Fig. 5A-D, Extended Data Fig. 5A**). However, the organotropism of metastatic gastric tumors had a genotype preference: mice harboring *Apc*-null GS tumors showed a higher frequency of liver metastasis (8/9, 88% of mice) compared to those harboring p53-null CIN (5/9, 55% of mice) or Msh2-null MSI (3/10, 30% of mice) tumors (**Fig. 5E**). Interestingly, the propensity of *Apc*-null GS tumors to colonize the liver was also noted following introduction of a subset of *Apc*-null tumor-derived lines following tail vein injection, an experimental metastasis assay that strongly favors seeding to the lung (**Extended Data Fig. 5B**). By stark contrast, mice harboring MSI tumors were markedly less capable of metastasizing to the lungs (30%, compared to 55% and 67% for GS and CIN tumors, respectively) (**Fig. 5F**).

**Figure 5.**
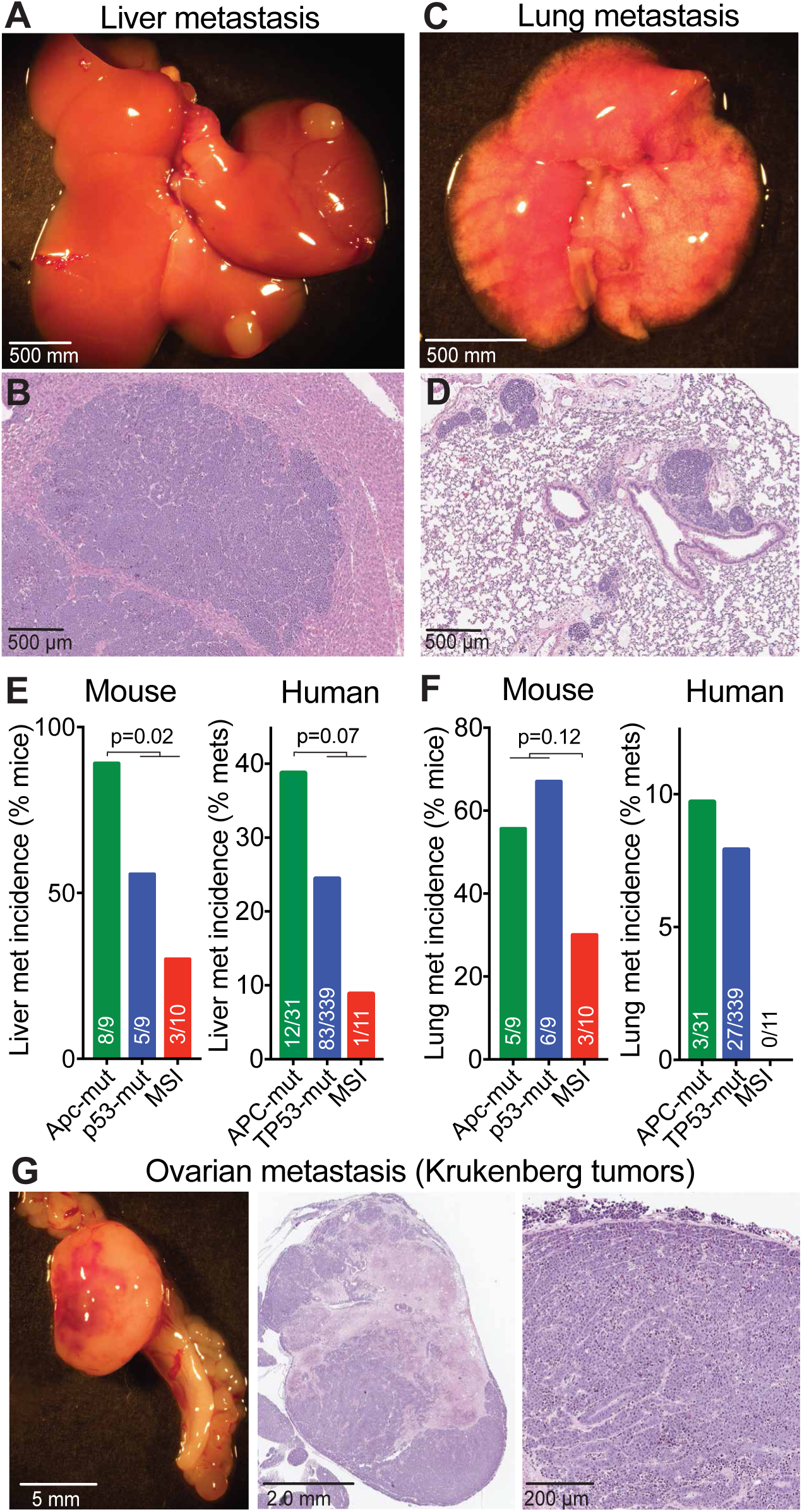
Gastric cancer EPO-GEMMs display metastatic patterns that recapitulate the human disease. (A-B) Representative macroscopic (A) and H&E-stained histological (B) images of liver metastases from a *MYC-p53^-/-^*gastric cancer EPO-GEMM. (C-D) Representative macroscopic (C) and H&E-stained histological (D) images of lung metastases from a *MYC-p53^-/-^* gastric cancer EPO-GEMM. (E-F) Incidence of liver (E) or lung metastases (F) in gastric cancer EPO-GEMMs of the indicated genotypes (left graphs), and among the MSK-IMPACT cohort of esophagogastric cancer patients with the corresponding genetic alterations. Exact number of independently analyzed tumors is indicated. Statistical analysis by Fisher’s exact test. (G) Representative macroscopic (left) and H&E-stained histological (middle and right) images of an ovarian metastasis from a *MYC-p53^-/-^* gastric cancer EPO-GEMM.

The different metastatic profiles of gastric cancer subtypes in mice were unexpected. To determine if similar patterns exist in the human disease, we took advantage of the clinical annotation of tumors in MSK-IMPACT to link tumor genotype to metastasis pattern in gastric cancer patients (**Fig. 5E-F**). Remarkably, of all metastatic samples that harbored *APC* mutations, 39% (12/31) were derived from the liver compared to only 24% (83/339) and 9% (1/11) of metastases harboring *TP53* or mismatch repair mutations, respectively (**Fig. 5E**). Likewise, none (0/11) of the mismatch repair mutant metastases were derived from the lung, in contrast to 10% (3/31) and 8% (27/339) of *APC*-and *TP53*-mutant metastases, respectively (**Fig. 5F**). Corroborating these results, WNT pathway alterations were associated with a significant increase in the incidence of liver but not lung metastasis, whereas mutations in *TP53* correlated with a small increase in the metastasis incidence to both organs (**Extended Data Fig. 5C**)^42^.

A subset of gastric cancer patients develops “Krukenberg tumors”^43, 44^, which arise from metastasis to the ovary. While poorly understood, these tumors are clinically important as they often arise in young women (median age 45 years) and confer a dismal prognosis^45^. Owing in part to the lack of model systems, the etiology of these tumors remains unresolved, with an ongoing debate about the lymphatic vs. hematogenous route of dissemination from the primary tumor^45^. Remarkably, a subset of gastric cancer EPO-GEMM animals developed metastases in the ovaries (**Fig. 5G**). Consistent with a hematogenous route of spread, the capacity for ovarian metastasis was maintained in a subset of primary tumor lines assayed by tail vein injection (**Extended Data Fig. 5D**). Together, these data highlight the relevance of EPO-GEMMs as a robust platform to study metastatic gastric cancer and reveal a role for tumor genotype in metastatic organotropism.

### NK cells suppress gastric cancer metastasis

Studies aimed at understanding mechanisms of metastasis have primarily focused on transcriptional and epigenetic changes within tumor cells. More recent work using carcinogen-induced or transplantation models has revealed a role for immune cells in both facilitating and limiting metastatic spread but little is known about the influence of the immune system on metastasis in autochthonous, genetically defined settings that are closest to the human scenario^46–48^. To address this gap, we harnessed the ability of the EPO-GEMM approach to engineer genetically defined tumors in different recipient strains. To this end, *MYC-Apc^-/-^*tumors were generated via stomach electroporation of: (1) wild-type C57BL/6 mice, which are fully immune-competent; or (2) Rag2-Il2rg double-knockout (R2G2) mice, which are deficient in T, B and NK cells, and have reduced levels of neutrophils, macrophages and dendritic cells. Tumor-bearing R2G2 immune deficient mice showed reduced survival and a greater incidence of liver metastasis compared to immune competent recipients (**Fig. 6A-C**). These data indicate that the immune system potently suppresses metastasis in an autochthonous gastric cancer model.

**Figure 6.**
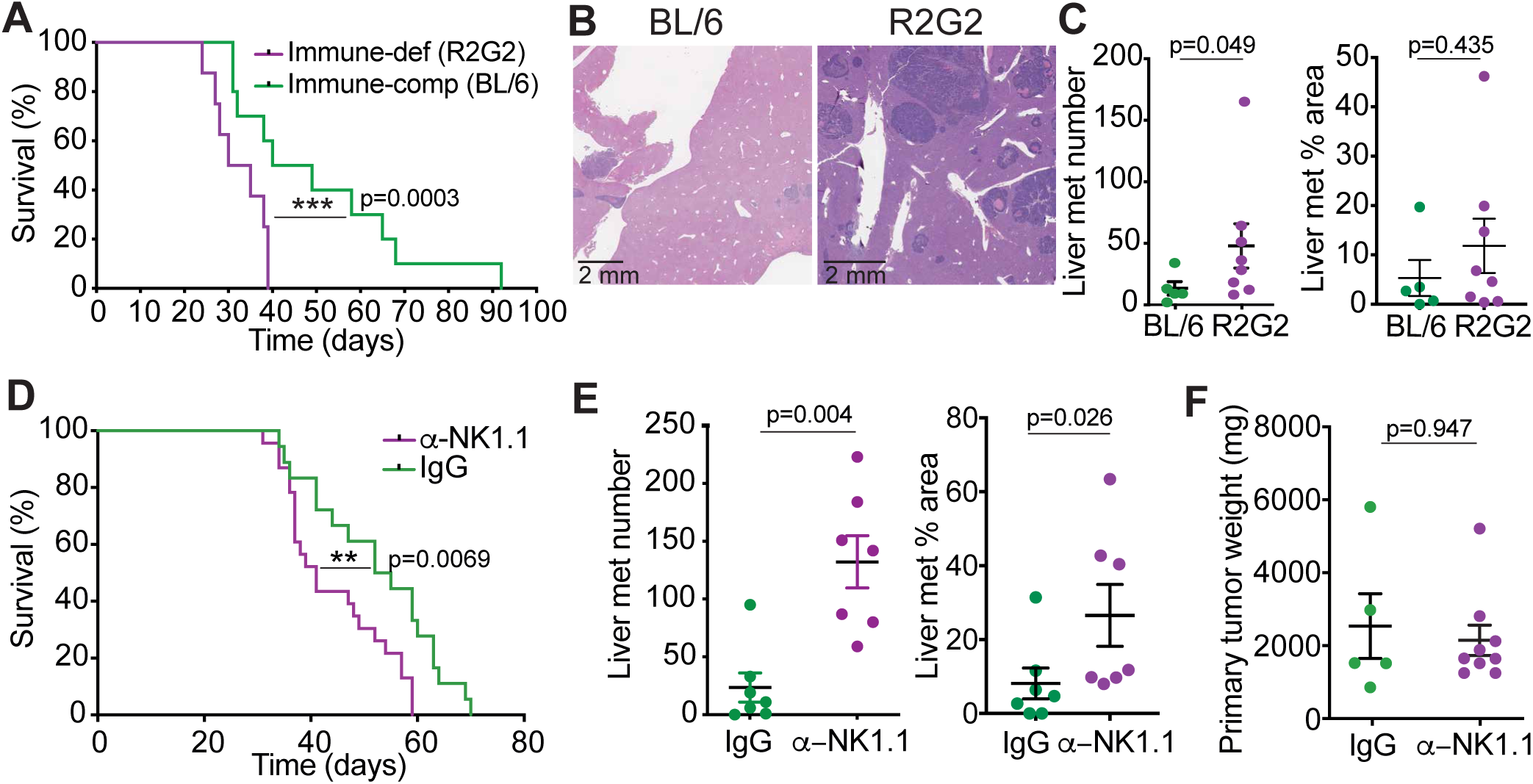
NK cells suppress gastric cancer metastasis. (A) Kaplan-Meier survival curve of immune-competent C57BL/6 (BL/6, green; same cohort as shown in Fig. 2C) or immune-deficient R2G2 (purple, n=8) mice with electroporation-induced *MYC-Apc^-/-^*gastric cancer. (B) Representative hematoxylin and eosin (H&E) staining of liver metastases from mice in (A). (C) Quantification of the number of liver metastases (left) and the percentage area of total liver occupied by metastases (right) from a subset of mice in (A) (BL/6 n=5; R2G2 n=8). (D) Kaplan-Meier survival curve of BL/6 *MYC-Apc^-/-^*gastric cancer EPO-GEMMs treated with an NK1.1-targeting antibody (purple, n=23) or IgG control (green, n=18). (E) Quantification of the number of liver metastases (left) and the percentage area of total liver occupied by the metastasis (right) from a subset of mice in (D). (IgG n=7; NK1.1 n=7) (F) Matching primary tumor weights from a subset of mice in (D). (IgG n=5; NK1.1 n=9) Statistical analysis: (A, D) Log-rank test, (C, E, F) Mann-Whitney test.

NK cells are a prominent immune cell type that can limit the extent of metastasis in certain experimental metastasis models^49^. To assess the role of NK cells in an autochthonous context, we administered NK1.1-targeting antibodies in immune-competent mice from the time of electroporation, which results in systemic depletion of NK and related cells^47^. As in the immune-deficient recipients, NK1.1-treated immune-competent mice displayed decreased overall survival and increased liver metastasis compared to isotype-treated controls (**Fig. 6D-E**). No difference in primary tumor size was observed at endpoint, suggesting that the contribution of NK cells to improving animal survival was mainly due to the suppression of metastasis (**Fig. 6F**). Reinforcing this point, similar results were observed when NK cell depletion was induced after detection of a palpable primary tumor (**Extended Data Fig. 6A-B**) or in experimental metastasis assays that examine metastatic potential of circulating tumor cells following tail vein or intrasplenic injection (**Extended Data Fig. 6C-F**). Therefore, NK cells play a critical role in curtailing gastric cancer metastasis.

### CD8^+^ T cells provide an added layer of metastasis surveillance in MSI gastric cancer

To further characterize how the immune system restricts metastatic spread across gastric cancer subtypes, we performed experimental metastasis assays using cell lines derived from primary EPO-GEMM tumors representing the GS (*MYC-Apc^-/-^*), CIN (*MYC-p53^-/-^*), and MSI (*MYC-p53^-/-^-Msh2^-/-^*) subtypes. Mice were treated with an NK1.1 vs. IgG control antibody twice per week starting two days before tail vein injection, and monitored for development of metastases. NK cell depletion led to a significant increase in both liver and lung metastatic burden in mice injected with either CIN or MSI cancer cells (**Fig. 7A**). However, the metastatic potential of MSI tumors remained lower than that of MSS tumors even following NK cell depletion, an effect that was particularly pronounced in the lung (**Fig. 7A**). Corroborating these results, only 17% (2/12) of immune-competent mice injected with MSI gastric cancer cells developed overt lung metastases, compared to 75% (8/12) of mice injected with MSS tumor cells (**Fig. 7B-C**). At the same time, both the MSI and MSS subtypes showed a similar ability to form lung metastases following tail vein injection into immunodeficient R2G2 mice, indicating there was no appreciable difference in their cell intrinsic potential to colonize the lung (**Fig. 7B-C**).

**Figure 7.**
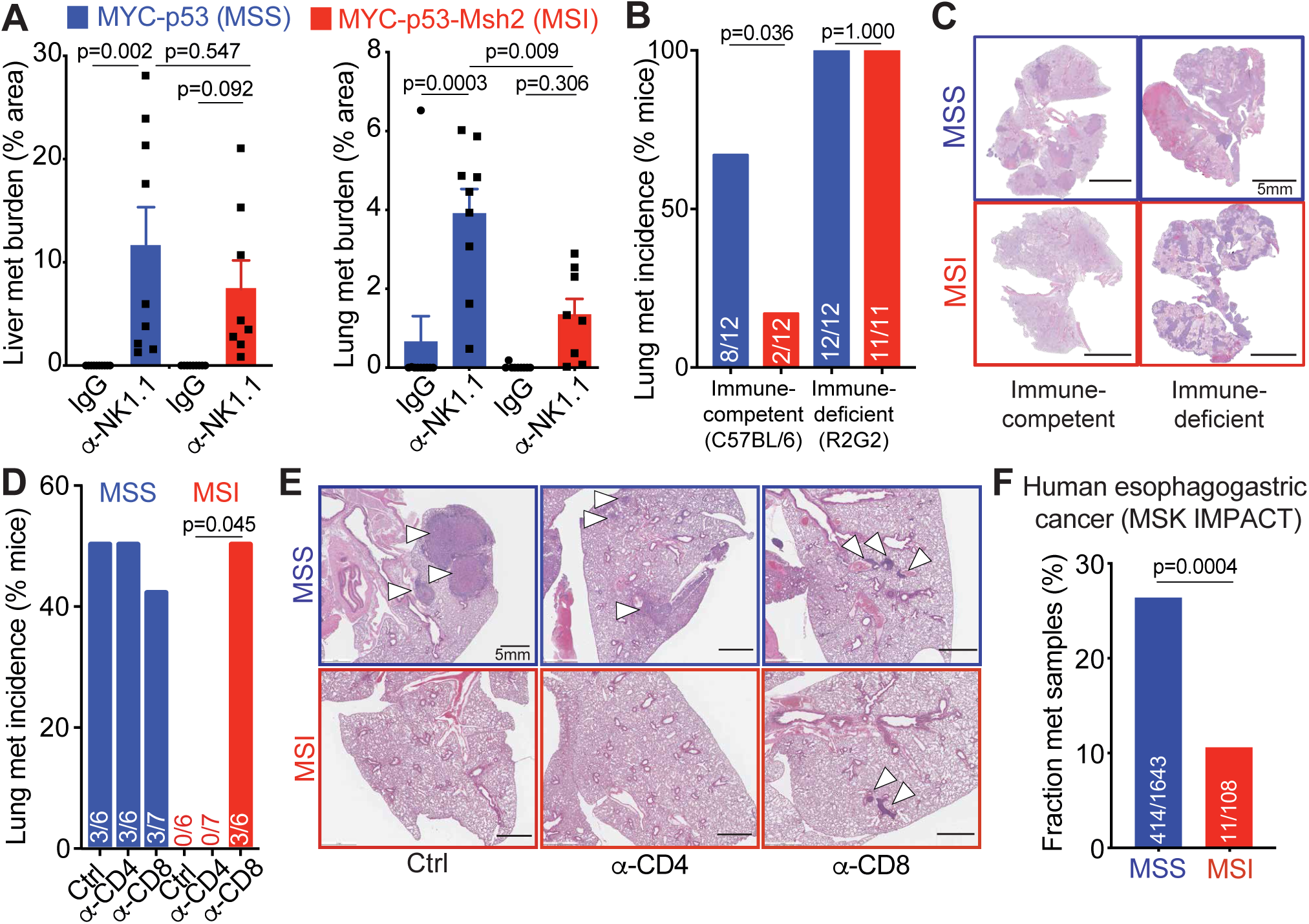
CD8+ T cells provide an added layer of metastasis immune surveillance in MSI tumors. (A) Metastatic burden (% tumor area) in the liver (left) or lung (right) of BL/6 mice after tail vein injection of *MYC-p53^-/-^* (MSS, blue, n=9-10) or *MYC-p53^-/-^-Msh2^-/-^* (MSI, red, n=8-9) gastric cancer cells. Mice were treated with either an NK1.1-targeting antibody or IgG control. (B) Incidence of lung metastasis after tail vein injection of MSS or MSI gastric cancer cells into immune-competent (C57BL/6) or immune-deficient (R2G2) mice. Exact numbers of independent mice are indicated on each bar. (C) Representative hematoxylin and eosin (H&E) staining of lungs isolated from mice in (B). (D) Incidence of lung metastasis after tail vein injection of MSS or MSI gastric cancer cells into immune-competent (C57BL/6) mice that were treated with either CD4- or CD8-targeting antibodies, or an IgG control. Exact numbers of independent mice are indicated on each bar. (E) Representative hematoxylin and eosin (H&E) images of lungs isolated from mice in (D). (F) Fraction of metastatic samples in the MSK-IMPACT cohort of esophagogastric cancer patients with either MSS or MSI disease. Exact number of independently analyzed tumors is indicated on each bar. Statistical analysis: (A) Ordinary one-way ANOVA, (B, D, F) Fisher’s exact test.

Since mismatch repair-deficient tumors can present high amounts of neoantigens on MHC molecules and elicit a T cell-mediated immune response (**Fig. 3C-F**)^13^, we reasoned that cytotoxic T cells may also contribute to the surveillance of MSI tumors. We therefore depleted either CD4^+^ or CD8^+^ cells in fully immune-competent hosts and assessed the metastatic potential of either MSS or MSI tumor cells following tail vein injection. Unlike MSS cells, which formed metastases across all conditions, MSI cells seeded metastatic tumors only in the CD8-depleted condition (**Fig. 7D-E**). Importantly, these results are consistent with reduced metastasis incidence that we observed in MSI patients (**Fig. 7F**). In sum, these data reveal a bi-modal surveillance of gastric cancer metastasis – a genotype-agnostic control by NK cells supplemented with MSI-specific control by CD8^+^ T cells – and support the use of the EPO-GEMM platform to study tumor-host interactions influencing metastatic spread.

## DISCUSSION

Here we present a suite of fully somatic mouse models of gastric cancer produced by delivery of genetic elements directly to the stomach using tissue electroporation that we refer to as gastric cancer EPO-GEMMs. By combining different mutational events associated with distinct tumor subtypes, we demonstrate that this platform can produce models of all three non-viral molecular subtypes of human gastric cancer. Besides sharing the underlying genotypes, these models mirror the defining histological and transcriptional properties of their respective human subtypes and present similar patterns of chromosomal (in)stability and mutational signatures. Perhaps most importantly, each model reproducibly metastasizes to clinically relevant anatomical sites. These features demonstrate the relevance of gastric cancer EPO-GEMMs for discovery and preclinical studies, including in the context of metastasis.

The genetic flexibility of the gastric cancer EPO-GEMMs eliminates the need for extensive strain intercrossing and enables rapid testing of any genetic combination by simply changing the sequence of electroporated constructs. Moreover, synchronous cohorts of animals that will develop genotypically defined tumors can be produced in a day, thereby greatly simplifying the execution of mechanistic and preclinical studies. As such, gastric cancer EPO-GEMMs offer advantages over carcinogen-induced models, which do not produce genetically defined tumors^50–52^, and Cre/lox-based models, which are limited to available germline strains, yield asynchronous cohorts, and produce substantial animal waste as unavoidable byproducts of strain intercrossing^22, 23^. Furthermore, EPO-GEMMs produce focal cancers in adult mice, avoiding the confounding effects of tissue-wide gene activation/inactivation during embryogenesis or, conversely, the requirement for tamoxifen (which can induce gastric metaplasia^53–55^) to recombine germline alleles later on. Finally, EPO-GEMMs offer the unique capability to readily change the host, which provides a flexible and robust platform to study tumor-host interactions in a manner that is impractical for traditional GEMMs.

We illustrate the power of EPO-GEMMs to uncover new biology by gaining novel insights into gastric cancer pathogenesis. As one example, by comparing the histopathology of each tumor subtype we noted that late-stage p53-null CIN tumors, which are predominantly moderately differentiated, harbor undifferentiated regions that lacked E-cadherin expression and resembled GS tumors produced by E-cadherin inactivation. These observations suggest that CIN and GS subtypes are subject to the forces of convergent evolution and, accordingly, we noted the occurrence of *TP53* and *CDH1* mutations are mutually exclusive in the human disease.

Other important insights arising from the initial characterization of these models relate to the nature and mechanisms of metastatic organotropism. First, a subset of EPO-GEMM animals develop ovarian tumors with features of Krukenberg tumors, an enigmatic but clinically relevant facet of gastric cancer presentation that has not been previously modeled. Our results demonstrate that these tumors can arise from different gastric cancer genotypes and establish hematogenous migration as a viable route of gastric cancer spread to the ovary. Second, EPO-GEMM models display a genotype-specific pattern of metastatic organotropism that, though not previously known, was mirrored in human patients. Hence, *Apc*-null GS tumors showed a preferential ability to metastasize to the liver (a pattern that may extend to other cancer types^25, 42^), whereas *Msh2*-deficient MSI tumors were poorly metastatic in general and showed a particularly pronounced impairment in lung metastasis. Finally, by targeting different recipient strains, we identified genotype-specific mechanisms of metastasis immune surveillance. While NK cells played a crucial role in suppressing metastatic spread in all non-viral molecular subtypes of gastric cancer^47, 49, 56–59^, MSI tumors were kept in check by an additional layer of immune surveillance provided by CD8^+^ T cells. This added layer of protection may explain the improved prognosis of MSI patients with gastric and other gastrointestinal cancers^60–62^.

Metastatic gastric cancer is a global health problem that is increasing in incidence. In many ways, the current state of gastric cancer research is comparable to where pancreatic cancer was two decades ago – a lethal cancer that is understudied, in part, due to the lack of faithful experimental models. The “KPC” mouse produced with conditional *Kras^G12D^* and *p53* mutant alleles revolutionized the study of pancreatic cancer and remains a gold standard model for studying the disease today^63^. With their flexibility and breadth, gastric cancer EPO-GEMM may have a similar impact, while at the same time enabling the study of a broader range of disease subtypes in reduced time and with less animal waste. Furthermore, as shown here, molecular studies on tumor-host interactions – now appreciated as central to cancer biology and therapy response – are straightforward. We anticipate that this platform will facilitate basic discovery efforts and accelerate the development of urgently needed therapeutic strategies for this deadly but understudied disease.

## METHODS

### Cell Lines and Compounds

The following cell lines were used in this study: *MP, MApc, MP.MSH,* which were derived from EPO-GEMM gastric tumors with these genotypes. To generate these cell lines, gastric tumors were minced, digested in DMEM media containing 3 mg/ml Dispase II (Gibco) and 1mg/ml Collagenase IV (C5138; Sigma) for 30 minutes at 37C, and plated on 10-cm culture dishes coated with 100 ug/ml collagen (PureCol; 5005; Advanced Biomatrix). Primary cultures were passaged at least three times to remove fibroblast contamination. Cells were maintained in a humidified incubator at 37C with 5%CO_2_ and grown in DMEM supplemented with 10% FBS and 100 IU/ml penicillin/streptomycin. All cell lines used were negative for mycoplasma.

### Reagents

For *in vivo* experiments mice were treated with anti CTLA4 (200 ug Bio X Cell; BE0131) 3 times per week per IP injection. Anti NK1.1 (250ug; Bio X Cell; BE0036), anti CD8 (200ug; Bio X Cell; BE0061), anti CD4 (200ug; Bio X Cell; BP00031) or the respective isotype control (Bio X Cell; BE0290; Bio X Cell; BE0090) was given twice per week by IP injection.

### Animal Studies

All mouse experiments were approved by the Memorial Sloan Kettering Cancer Center Institutional Animal Care and Use Committee. All relevant animal use guidelines and ethical regulations were followed. Mice were maintained under specific pathogen-free conditions, and food and water were provided ad libitum. The following mice were used: C57BL/6N background, *Nu/Nu* Nude mice (purchased from The Jackson laboratory) and Rag2-Il2rg double knockout mice (R2G2). Mice were female and/or male and were used at 8-12 weeks of age and were kept in group housing. Mice were randomly assigned to the experimental groups.

### EPO-GEMMs

8- to 12-week-old WT C57BL/6 mice were starved for two hours prior to the procedure. Mice were anesthetized with isoflurane and the surgical site (epigastrium) scrubbed with a povidone-iodine scrub (Betadine) and rinsed with 70% alcohol. After opening the peritoneal cavity, the stomach was mobilized and opened up in the area of the forestomach. Afterwards the inside of the stomach was rinsed with saline to remove any residual food. Subsequently, the plasmid mix (50ul; see specifications below) was injected into the epithelial compartment in the corpus/antrum region using a 30-gauge syringe and tweezer electrodes were tightly placed around the injection bubble. Two pulses of electrical current (75V) given for 35-milisecond lengths at 500-milisecond intervals were then applied using an *in vivo* electroporator (NepaGene NEPA21 Type II Electroporator). After electroporation, the stomach was closed with absorbable sutures and the peritoneal cavity was rinsed with 0.5 mL of prewarmed saline. The peritoneal cavity was sutured, and the skin closed with skin staples. The mice were kept at 37°C until awoke, and post-surgery pain management was done with injections of buprenorphine and/or meloxicam for the 3 following days. Tumor formation was assessed by palpation or ultrasound imaging, and mice were sacrificed following early tumor development or at endpoint. Genome editing in EPO-GEMM tumors was confirmed by Sanger sequencing.

To generate EPO-GEMM tumors in C57BL/6 WT mice, the following vectors and concentrations were used: a pT3-MYC transposon vector (5 μg), a Sleeping Beauty transposase (SB13; 1 μg), and/or a pX330 CRISPR/Cas9 vector (20 μg; Addgene #42230) targeting the respective tumor suppressor genes. The Sleeping Beauty transposase (SB13) and the pT3 transposon vector were a generous gift from Dr. Xin Chen (University of California, San Francisco, San Francisco, CA). The pX330 vector was a gift from Feng Zhang of Broad Institute (Addgene plasmid # 42230).

The following sgRNAs were used to target the respective tumor suppressor gene locus:

p53: ACCCTGTCACCGAGACCCC

APC: GCAGGAACCTCATCAAAACG

CDH1: CCCGTTGGCGTTTTCATCAT

MSH2: GACAAAGATTGGTTAACCAG

To generate the pX330 vector containing two sgRNAs, the vector was opened using the *Xba*I cloning site and the sgRNA-casette containing the second guide was PCR cloned into the vector using the following primers: XbaI U6 forward: ATGCTTCTAGAGAGGGCCTATTTCCCATGATT and NheI gRNA scaffold reverse: ATGTCGCTAGCTCTAGCTCTAAAACAAAAAAGC.

### Experimental Metastasis Assays

1 x 10^5^ *MP*, *MPMsh2* or *MApc* gastric tumor cells were resuspended in 400 μL of PBS and tail vein injected into 8- to 12-week-old C57BL/6N, Nude or R2G2 mice.

### Analysis of Metastasis Burden

The presence of lung, and liver metastases were determined at experimental endpoint by gross examination under a dissecting scope. Metastasis burden and the total number of individual metastases was further quantified from hematoxylin and eosin (H&E)–stained sections.

### Histological Analysis

Tissues were fixed overnight in 10% formalin, embedded in paraffin, and cut into 5 μm sections. Sections were subjected to hematoxylin and eosin (H&E) staining. Immunohistochemical and immunofluorescence stainings were performed following standard protocols. The following primary antibodies were used: E-Cadherin (1:500,BD Bioscience, 610181), H+K (1:1000, MBL International Corporration, D032-3), Ki67 (1:100, Abcam, AB16667), CK8 (1:1000, BioLegend, 904801), MSH2 (1:200, Cell Signaling, D24B5), MYC (1:100, Abcam, AB32072), and Vimentin (1:200, Cell Signaling, 5741).

### Flow Cytometry

For *in vivo* sample preparation, gastric tumors were processed into small pieces, digested in RPMI containing 2mg/ml Collagenase D and 100ug/ml DNase I for 30 minutes at 37C, filtered through a 70μm strainer, washed with PBS, and red blood cell lysis was achieved with an ACK (Ammonium-Chloride-Potassium) lysing buffer (Lonza). Cells were washed with PBS, resuspended in FACS buffer and used for subsequent analysis. The following fluorophore-conjugated antibodies were used (‘m’ prefix denotes anti-mouse): m.CD45 (AF700), m.CD3 (PE-Cy7), CD3 (AF488), CD4 (BUV395), CD8 (PECy7), CD11c (BV650), m.CD3 (BV650), m.CD4 (BUV737), m.CD8 (FITC), m.CD11c (BV785). Flow cytometry was performed on a LSRFortessa instrument (BD Biosciences) and data were analyzed using FlowJo (TreeStar).

### RNA Extraction, RNA-seq Library Preparation and Sequencing

Total RNA was isolated from: MP, MP.MSH2, MAPC and MCDH1 tumors. Library preparation and sequencing were performed at the Integrated Genomics Operation (IGO) Core at MSKCC. RNA-seq libraries were prepared from total RNA. After RiboGreen quantification and quality control by Agilent BioAnalyzer, 100-500ng of total RNA underwent polyA selection and TruSeq library preparation according to instructions provided by Illumina (TruSeq Stranded mRNA LT Kit, RS-122-2102), with 8 cycles of PCR. Samples were barcoded and run on a HiSeq 4000 or HiSeq 2500 in a 50bp/50bp paired end run, using the HiSeq 3000/4000 SBS Kit or TruSeq SBS Kit v4 (Illumina).

### RNA-seq Read Mapping, Differential Gene Expression Analysis and Heatmap Visualization

RNA-Seq data was analyzed by removing adaptor sequences using Trimmomatic^64^. RNA-Seq reads were then aligned to GRCm38.91 (mm10) with STAR^65^ and transcript count was quantified using featureCounts^66^ to generate raw count matrix. Differential gene expression analysis and adjustment for multiple comparisons were performed using DESeq2 package^67^ between experimental conditions, using more than 2 independent biological replicates per condition, implemented in R (http://cran.r-project.org/). Differentially expressed genes (DEGs) were determined by > 2-fold change in gene expression with adjusted P-value < 0.05. For heatmap visualization of DEGs, samples were z-score normalized and plotted using pheatmap package in R.

### Functional Annotation of Gene Sets

Pathway enrichment analysis was performed in the resulting gene clusters with the Reactome database using enrichR^68^. Significance of the tests was assessed using combined score, described as c = log(p) * z, where c is the combined score, p is Fisher’s exact test p-value, and z is z-score for deviation from expected rank.

### Gene Set Enrichment Analysis (GSEA)

GSEA^69^ was performed using the GSEAPreranked tool for conducting gene set enrichment analysis of data derived from RNA-seq experiments (version 2.07) against signatures in the MSigDB database (http://software.broadinstitute.org/gsea/msigdb). The metric scores were calculated using the sign of the fold change multiplied by the inverse of the p-value.

### Gene Signature Score and Immune Cell Type Abundance Estimation

Rank-based single-sample gene set scoring method was calculated using package singscore in R^70^. Immune cell abundance estimation was based on LM22 signature^40^, which contains 547 gene signature matrix from 22 human immune cell types. LM22 signature and singscore were used to estimate gene expression profiles for each LM22 cell type.

### CNA Analysis

CNAs were inferred from sparse whole-genome sequencing data as described previously^71, 72^. In brief, 1 μg of bulk genomic DNA was extracted from gastric tumors using the DNeasy Blood and Tissue Kit (Qiagen) and sonicated using the Covaris instrument. Sonicated DNA was subsequently end-repaired/A-tailed, followed by ligation of TruSeq dual indexed adaptors. Indexed libraries were enriched via PCR and sequenced in multiplex fashion using the Illumina HiSeq2500 Instrument to achieve roughly 1 × 10^6^ uniquely mappable reads per sample, a read count sufficient to allow copy-number inference to a resolution of approximately 400 kb. For data analysis, uniquely mapped reads were counted in genomic bins corrected for mappability. Read counts were subsequently corrected for guanine cytosine content, normalized, and segmented using circular binary segmentation. Segmented copy-number calls are illustrated as relative gains and losses to the median copy number of the entire genome. Broad events (chromosome-wide and several megabase-sized events) are discernible in a genome-wide manner.

### Whole-Exome Sequencing (WES)

1 μg of bulk genomic DNA was extracted from gastric tumors using the DNeasy Blood and Tissue Kit (Qiagen) and WES was conducted and sequenced by BGI. The data was then processed through the Illumina (HiSeq) Exome Variant Detection Pipeline for detecting variants by the Bioinformatics Core at MSKCC. First, the FASTQ files were processed to remove any adaptor sequences at the end of the reads using cutadapt (v1.6). The files were then mapped using the BWA mapper (bwa mem v0.7.12). After mapping the SAM files were sorted and read group tags added using the PICARD tools. After sorting in coordinate order the BAMs were processed with PICARD MarkDuplicates. The marked BAM files were then processed using the GATK toolkit (v 3.2) according to the best practices for tumor normal pairs. They were first realigned using ABRA (v 0.92) and then the base quality values recalibrated with the BaseQRecalibrator. Somatic variants were then called in the processed BAMs using muTect (v1.1.7) for SNV and the Haplotype caller from GATK with a custom post-processing script to call somatic indels. Based on the information provided by Agilent SureSelect XT Mouse All Exon Kit, the total exome coverage was about 49.6MB. This coverage length was used to calculate mutations per MB and compared with publicly available mutational data downloaded from ^31^.

### Human Clinical Data Analysis

For transcriptomic analysis, TCGA Stomach Adenocarcinoma (STAD) RNA-seq data were downloaded through R package TCGAbiolinks^73^ to retrieve molecular subtypes, raw and normalized (TPM) count table. Patients with matched normal and tumor samples were identified and used to run subtype-specific differential expression analysis. Results were used to calculate the rank score for GSEA analysis and compare to EPO-GEMMs. Microarray data from GSE62254 were downloaded and processed through R package limma^74^. Differentially expressed genes between different molecular subtypes were identified and used for GSEA analysis. Normalized enriched scores (NES) were plotted and compared to EPO-GEMMs.

For metastasis analysis, human datasets were obtained through either the MSK Clinical Sequencing Cohort (MSK-IMPACT) via cBioPortal^75, 76^, or the MSK-MET cohort^42^, as indicated in the text. For the liver/lung tropism analysis (Fig. 5E-F), MSK-IMPACT samples were selected as follows: (1) Cancer Type: Esophagogastric Cancer, (2) Sample Type: Metastasis, and (3) Genotype (MUT: APC, MUT: TP53, or MSI_TYPE: Instable). Then, the fraction of selected samples that were located in the liver or lung was calculated as a percentage of all metastatic sites. For the MSS vs. MSI metastasis analysis (Fig. 7F), MSK-IMPACT samples were selected as follows: (1) Cancer Type: Esophagogastric Cancer, (2) MSI_TYPE: Stable or Instable. Then, the fraction of selected samples that were derived from metastatic sites was calculated as percentage of all (primary + metastatic) samples. For the liver/lung metastasis incidence analysis from the MSK-MET cohort (Extended Data Fig. 5C), Stomach Adenocarcinoma patients were filtered by the presence of WNT pathway or *TP53* mutations, and then analyzed for the incidence of liver or lung metastases, as described in the published study^42^. Statistical comparisons were performed through contingency table analyses using Fisher’s exact test in Prism 7.0 (GraphPad Software) for the MSK-IMPACT cohort, or as described previously for the MSK-MET cohort^42^.

### Immunoblotting

Cell lysis was performed using RIPA Buffer (Cell signaling Technology) supplemented with phosphatase inhibitors (5 mmol/L sodium fluoride, 1 mmol/L sodium orthovanadate, 1 mmol/L sodium pyrophosphate, and 1 mmol/L β-glycerophosphate) and protease inhibitors (Protease Inhibitor Cocktail Tablets, Roche). Protein concentration was determined using a Bradford Protein Assay Kit (Bio-Rad). Proteins were separated by SDS-PAGE and transferred to polyvinyl difluoride membranes (Millipore) according to the standard protocols. Membranes were immunoblotted overnight at 4C with antibodies against MSH2 (Cell Signaling, D24B5) or β-actin (Cell Signaling, 4970) in 5% BSA in TBS-blocking buffer. Membranes were incubated with secondary anti rabbit antibody (Cell Signaling, 7074) for 1h at room temperature. Blots were developed in Perkin-Elmer’s Western Lightning ECL per manufacturer’s instructions.

### Statistical Analysis and Figure Preparation

Data are presented as mean ± s.e.m. Statistical analysis was performed by Student’s t-test, ANOVA, Mann-Whitney test, Wilcoxon signed-rank test, or Fisher’s exact test using Prism 6.0 or 7.0 (GraphPad Software), as indicated in the respective figure legends. P-values <0.05 were considered to be statistically significant. Survival was determined using the Kaplan-Meier method, with log-rank test used to determine statistical significance. No statistical method was used to predetermine sample size in animal studies. Animals were allocated at random to treatment groups. Figures were prepared using Biorender for scientific illustrations and Illustrator CC 2021 (Adobe).

## DATA AVAILABILITY

RNA-seq data has been deposited in the Gene Expression Omnibus under GEO ID PRJNA818675. Source data provided with the paper. All other data supporting the findings of this study will be made available upon reasonable request to the corresponding authors.

## ACKNOWLEDGEMENTS

We thank A. Kulick, E. de Stanchina, Leah Zamechek and C. Zhu for technical assistance; N. Socci and the Bioinformatics core for assistance with whole-exome sequencing analysis and members of the Lowe laboratory for insightful discussions. This work was supported by a Memorial Sloan Kettering Cancer Center Support grant (P30 CA008748), Stand up to Cancer (SU2C) and R01CA233944-02 to S.W.L laboratory. J.L. was supported by the German Research Foundation (DFG) under Germany’s excellence strategy (EXC 2180 – 390900677) and a Shulamit Katzman Endowed Postdoctoral Research Fellowship. C.A. was supported by a postgraduate fellowship from La Caixa foundation and is the recipient of the Harold E. Varmus graduate student fellowship from the Gerstner Sloan Kettering graduate school. K.M.T. was supported by the Jane Coffin Childs Memorial Fund for Medical Research and a Shulamit Katzman Endowed Postdoctoral Research Fellowship. F.J.S.R. was supported by a Hanna Grey Fellowship from the Howard Hughes Medical Institute. J.F. was supported by the Care-for-Rare Foundation and the German Research Foundation (DFG) under Germany’s excellence strategy (EXC 2180 – 390900677). T.B. received support from the William C. and Joyce C. O’Neil Charitable Trust and the Memorial Sloan Kettering Single Cell Sequencing Initiative. S.W.L. is the Geoffrey Beene Chair of Cancer Biology and a Howard Hughes Medical Institute Investigator. We thank the following MSKCC core facilities for support: Integrated Genomics Operations, Flow Cytometry, Research Animal Resource Center, and Anti-tumor Assessment.

---

**Extended Data Figure 1.**
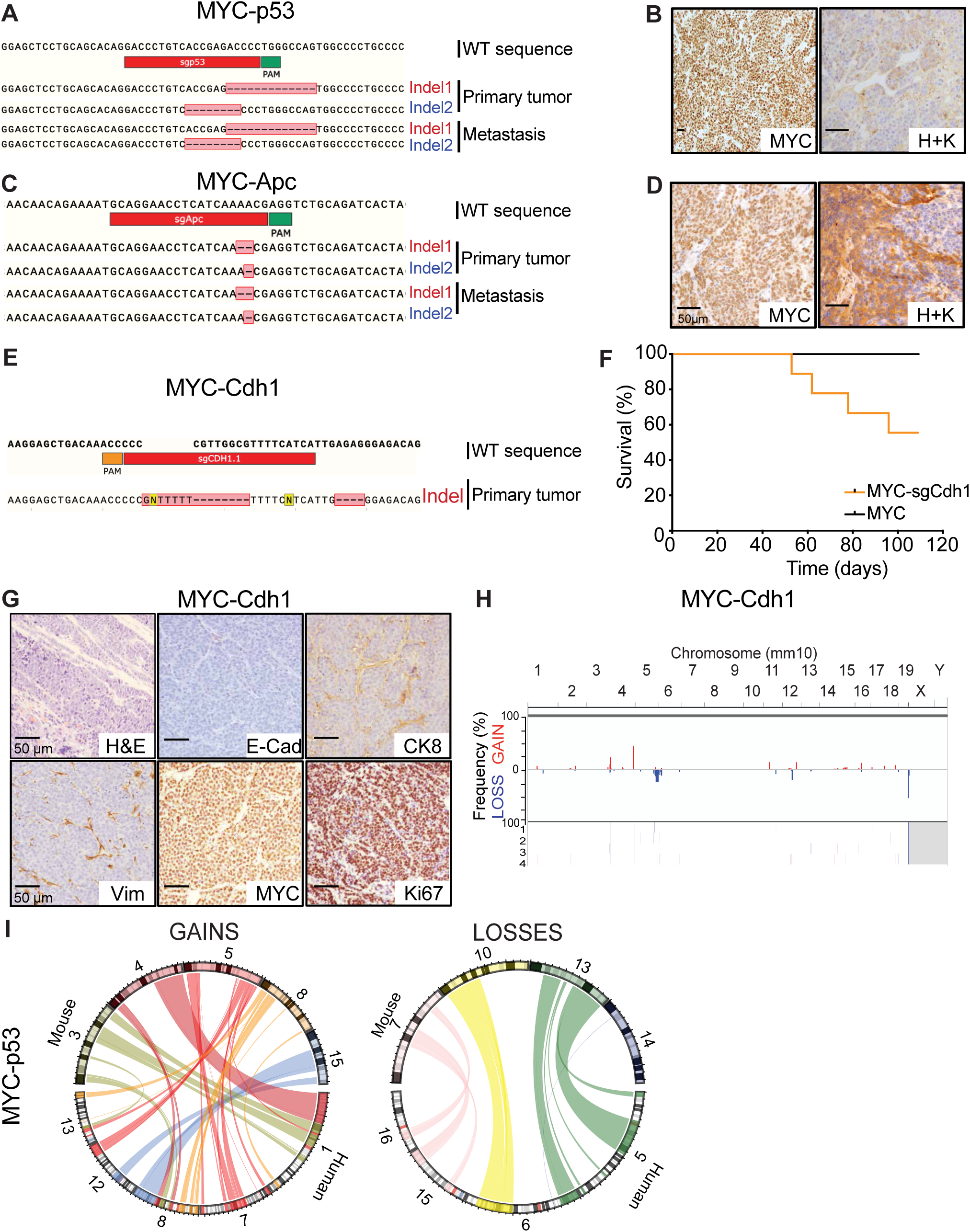
(A) Sanger sequencing results confirming editing of the respective gene locus targeted by the indicated CRISPR/Cas9-sgRNA in a MYC-p53 EPO-GEMM gastric tumor. (B) Representative immunohistochemistry staining for MYC and Hydrogen/Potassium ATPase (H+K) of a MYC-p53 EPO-GEMM gastric tumor. (C) Sanger sequencing results confirming editing of the respective gene locus targeted by the indicated CRISPR/Cas9-sgRNA in a MYC-Apc EPO-GEMM gastric tumor. (D) Representative immunohistochemistry staining for MYC and Hydrogen/Potassium ATPase (H+K) of a MYC-Apc EPO-GEMM gastric tumor. (E) Sanger sequencing results confirming editing of the respective gene locus targeted by the indicated CRISPR/Cas9-sgRNA in a MYC-Cdh1 EPO-GEMM gastric tumor. (F) Kaplan-Meier survival curve of C57BL/6 EPO-GEMMs with either *MYC-Cdh1^-/-^* (orange, n=9) or *MYC* only (black, n=4) gastric cancer. (G) Representative H&E and immunohistochemistry staining for E-Cadherin (E-cad), cytokeratin 8 (CK8), vimentin (Vim), MYC and Ki67 of a MYC-Cdh1 EPO-GEMM gastric tumor. (H) sWGS analysis of copy number alterations in MYC-Cdh1 (n=4) gastric EPO-GEMM tumors. Frequency plot is shown on the top and individual sample tracks are provided on the bottom. (I) Human-mouse synteny circos plots of recurrent copy-number gains and losses in MYC-p53 EPO-GEMM tumors.

**Extended Data Figure 2.**
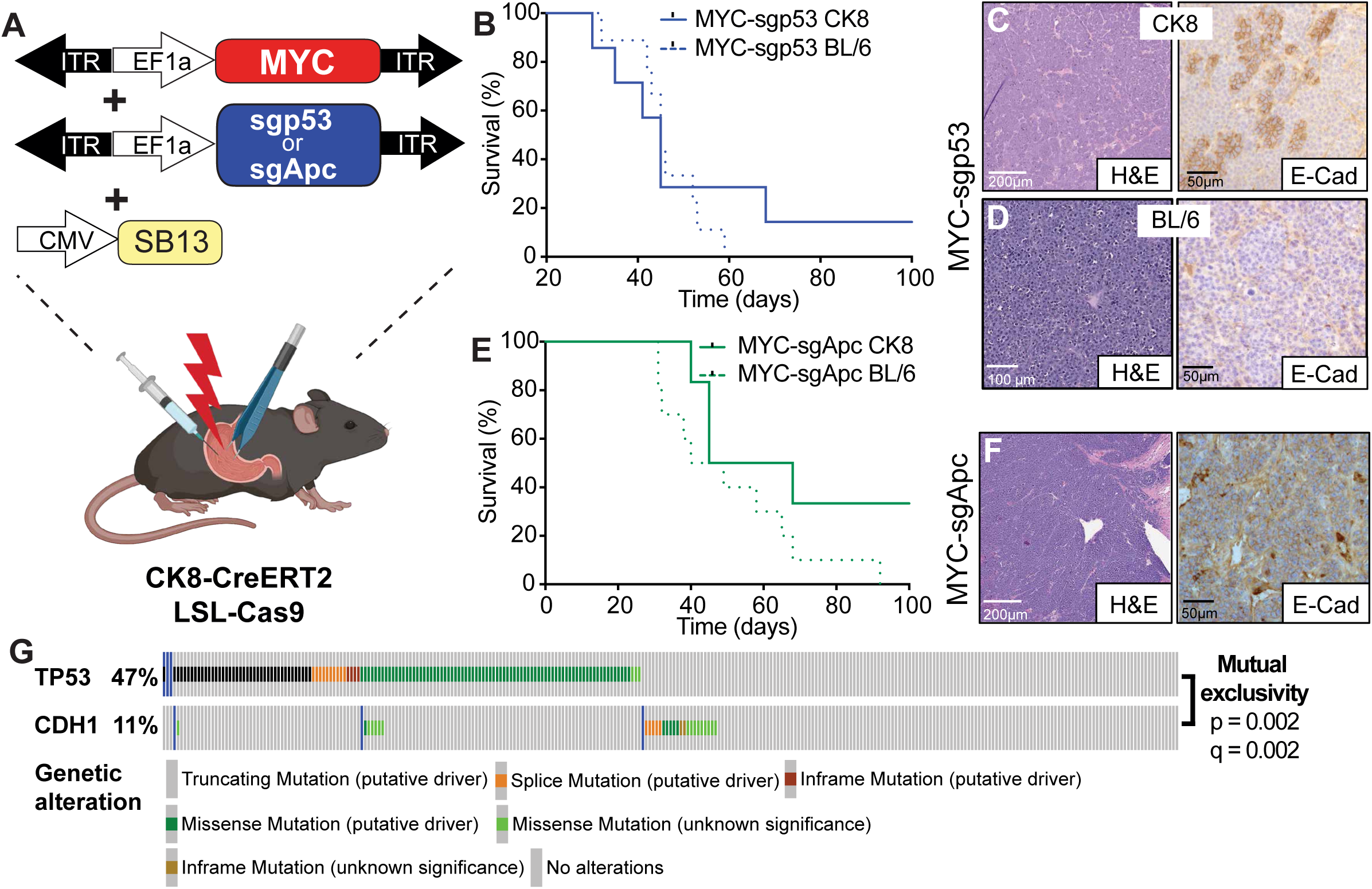
(A) Schematic of CK8-Cre restricted EPO-GEMM experiments. A transposon vector harboring an oncogene in combination with a Sleeping Beauty transposase (SB13) and/or a CRISPR-Cas9 vector targeting either *p53* or *Apc* were delivered into the stomach of CK8-CreERT2; LSL-Cas9 mice by direct *in vivo* electroporation. (B) Kaplan-Meier survival curve of C57BL/6 *MYC-p53^-/-^*EPO-GEMMs (blue dashed line, n=9, same cohort as shown in Fig. 2A) or CK8-CreERT2; LSL-Cas9 *MYC-p53^-/-^* EPO-GEMMs (blue line, n=7). (C) Representative H&E and immunohistochemistry for E-cadherin (E-Cad) in a *MYC-p53^-/-^* EPO-GEMM gastric tumor in a CK8-CreERT2; LSL-Cas9 mouse. (D) Representative H&E and immunohistochemistry for E-cadherin (E-Cad) of an undifferentiated *MYC-p53^-/-^* EPO-GEMM gastric tumor in a C57BL/6 (BL/6) mouse. (E) Kaplan-Meier survival curve of C57BL/6 *MYC-Apc^-/-^* EPO-GEMMs (green dashed line, n=10, same cohort as shown in Fig. 2C) or CK8-CreERT2; LSL-Cas9 *MYC-Apc^-/-^* EPO-GEMMs (green line, n=6). (F) Representative H&E and immunohistochemistry for E-cadherin (E-Cad) of a CK8-CreERT2; LSL-Cas9 *MYC-Apc^-/-^*EPO-GEMM gastric tumor. (G) MSK-IMPACT oncoprint displaying the genomic status of alterations in *TP53* and *CDH1* in gastric cancer patients. Alterations in *P53* and *CDH1* are mutually exclusive in this setting. Statistical analysis via cBioPortal^75, 76^.

**Extended Data Figure 3.**
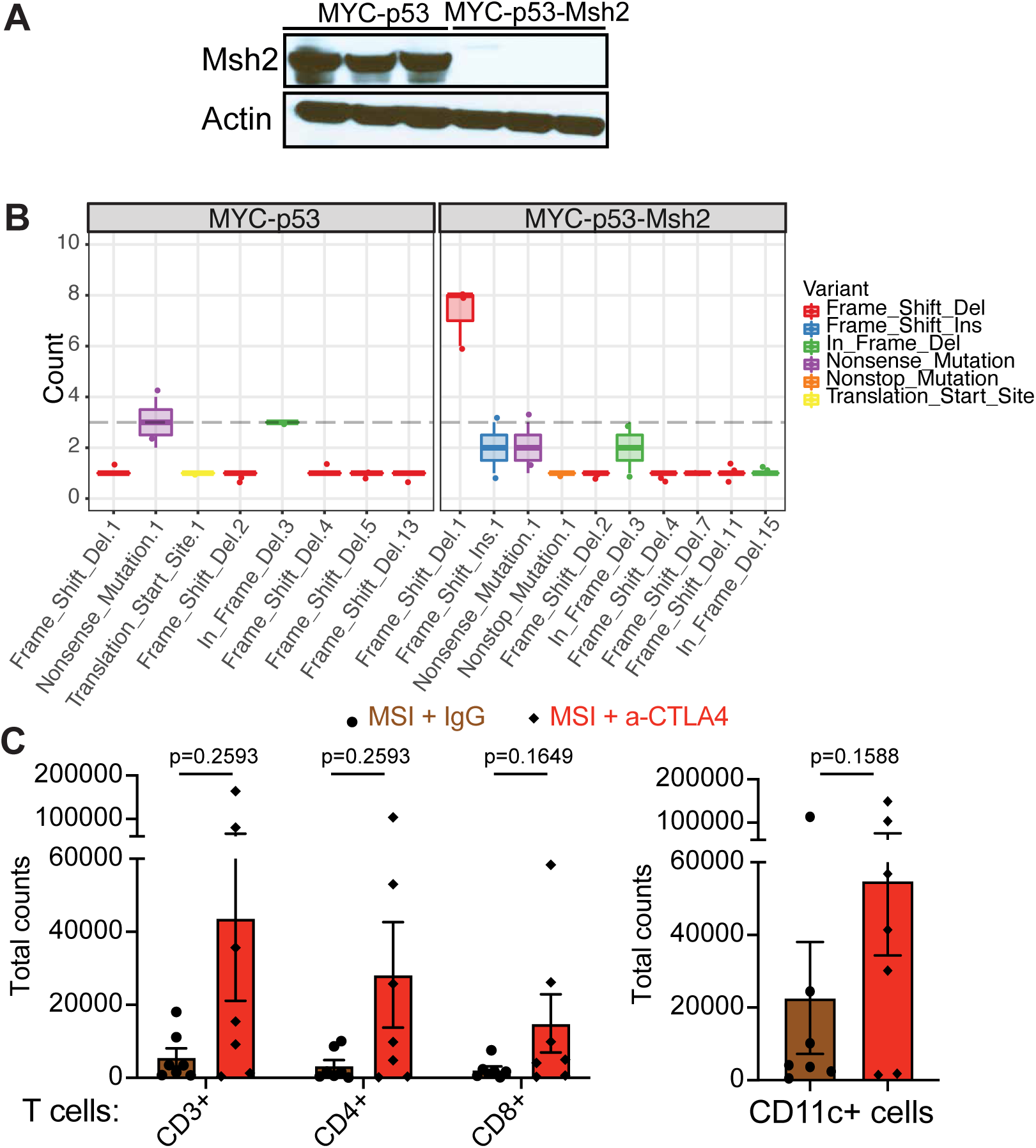
(A) Immunoblot of MSH2 and Actin (loading control) in MSI or MSS gastric cancer cell lines. (B) WES analysis of insertions (INS) or deletions (DEL) in either *MYC-p53^-/-^* or *MYC-p53^-/-^-Msh2^-/-^* gastric tumors (n=3 independent mice each). (C) Number of CD3+, CD4+ and CD8+ T cells (left) or CD11c+ cells (right) in MSI gastric tumors after treatment of mice with antibodies targeting CTLA-4 or IgG control (n=7 independent mice each). Statistical analysis by Mann-Whitney test.

**Extended Data Figure 4.**
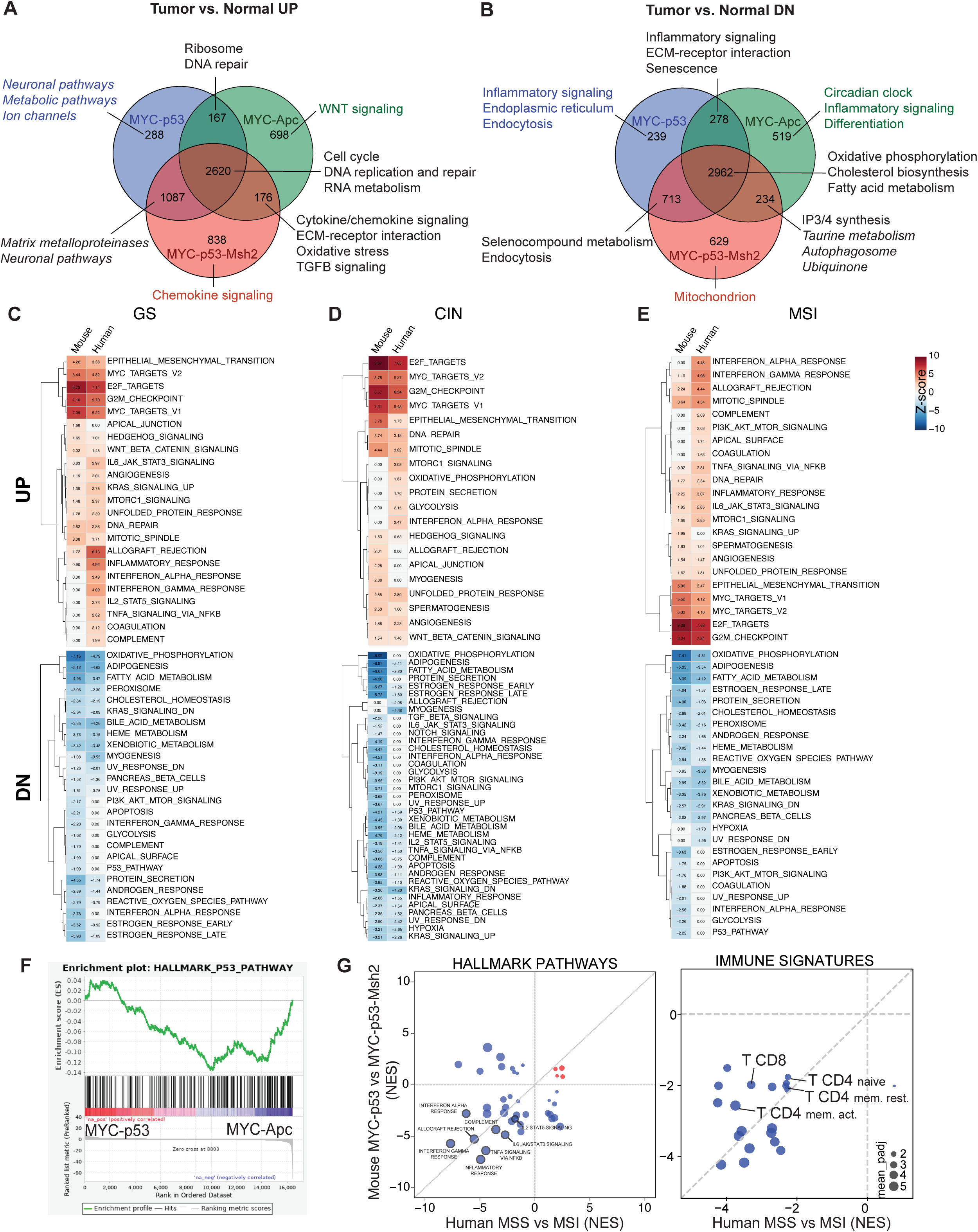
(A-B) Venn diagrams showing overlap of differentially up-(A) or down-regulated (B) genes (vs. normal stomach) for the indicated EPO-GEMM tumor genotypes. Key pathways enriched in each gene subset are labeled accordingly. Complete lists of pathway predictions are provided in **Extended Data Table 2**. (C-E) Complete lists of the Hallmark Pathways and NES scores shown in Figure 4B. (F) Gene set enrichment analysis (GSEA) comparing *MYC-p53^-/-^* and *MYC-Apc^-/-^* gastric tumors. (G) Comparison of GSEA NES scores for hallmark pathways (left) or Immune populations (right) enriched in mouse (y-axis) and human (x-axis) MSI gastric tumors. Highlighted are key immune populations enriched in MSI tumors. Circle size represents adjusted p-value. A complete list of NES scores is provided in **Extended Data Table 2**.

**Extended Data Figure 5.**
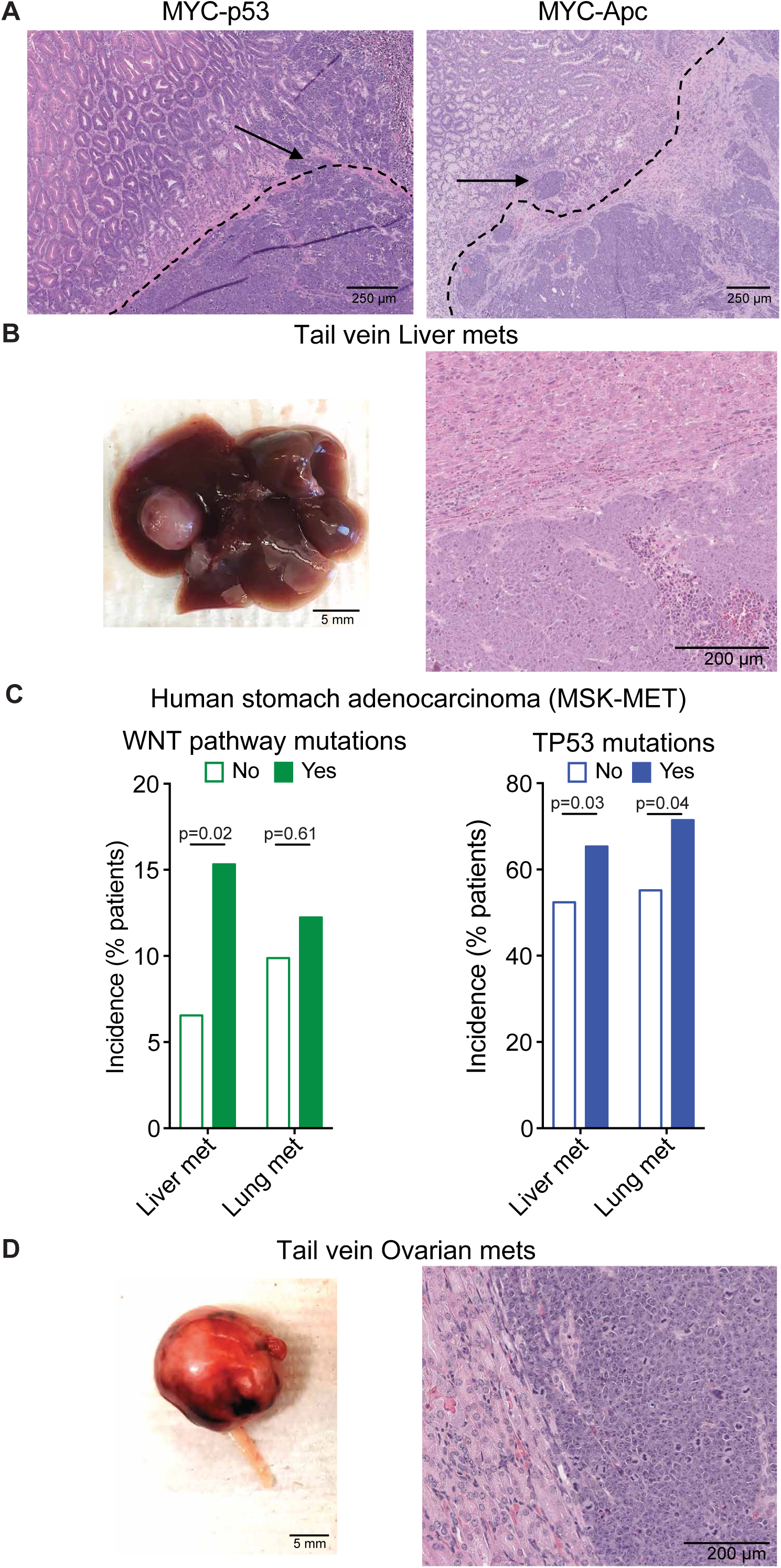
(A) H&E showing the delineation of gastric tumors within the normal stomach histology in C57BL/6 EPO-GEMMs with *MYC-p53^-/-^* (left) and *MYC-Apc^-/-^* (right). (B) Macroscopic (left) and H&E histology image (right) of liver metastasis in mice tail vein injected with a *MYC-p53^-/-^* gastric cancer cell line. (C) Incidence of liver and lung metastases among the MSK-MET cohort of gastric cancer patients with WNT pathway mutations or *TP53* mutations. Statistical analysis as described before^42^. (D) Macroscopic (left) and H&E histology image (right) of ovarian metastasis in mice tail vein injected with a *MYC-p53^-/-^* gastric cancer cell line.

**Extended Data Figure 6.**
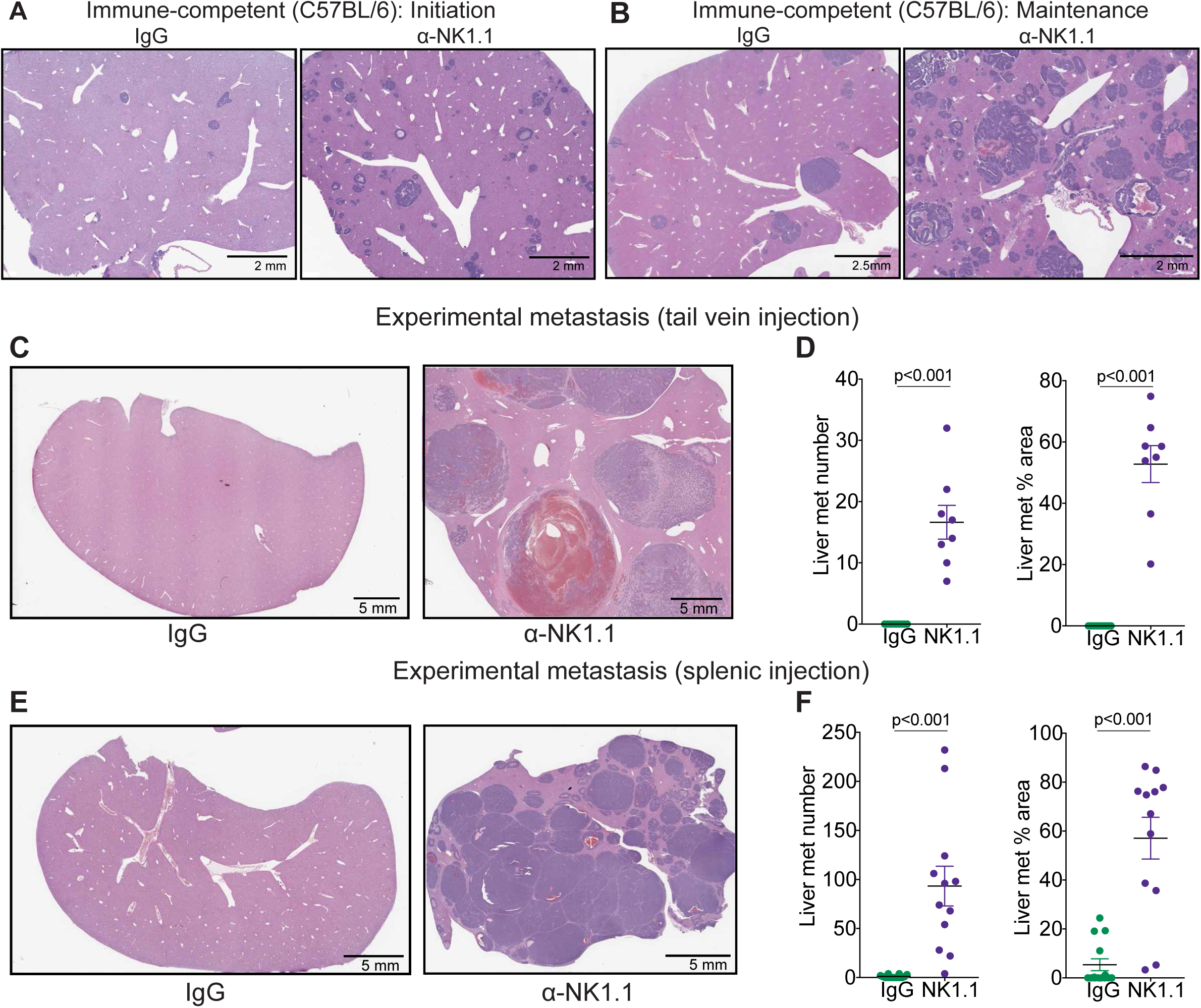
(A-B) Representative hematoxylin and eosin (H&E) staining of liver metastases of C57BL/6 *MYC-Apc^-/-^* EPO-GEMMs treated with an NK1.1-targeting antibody or the respective IgG control directly before tumor initiation (A) or after palpable tumor formation (B). (C) Representative hematoxylin and eosin (H&E) staining of livers of C57BL/6 mice after tail vein injection of *MYC-Apc^-/-^* gastric cancer cells and treatment with either an antibody targeting NK1.1 (right) or IgG control (left). (D) Quantification of the number of liver metastases (left) and the percentage area of total liver occupied by the metastasis (right) from mice in (C) (n=8 independent mice). Statistical analysis by Mann-Whitney test. (E) Representative hematoxylin and eosin (H&E) staining of livers of C57BL/6 mice after splenic injection of *MYC-Apc^-/-^* gastric cancer cells and treatment with either an antibody targeting NK1.1 (right) or IgG control (left). (F) Quantification of the number of liver metastases (left) and the percentage area of total liver occupied by the metastasis (right) from mice in (E) (n=12 independent mice). Statistical analysis by Mann-Whitney test.

**Extended Data Figure 7.**
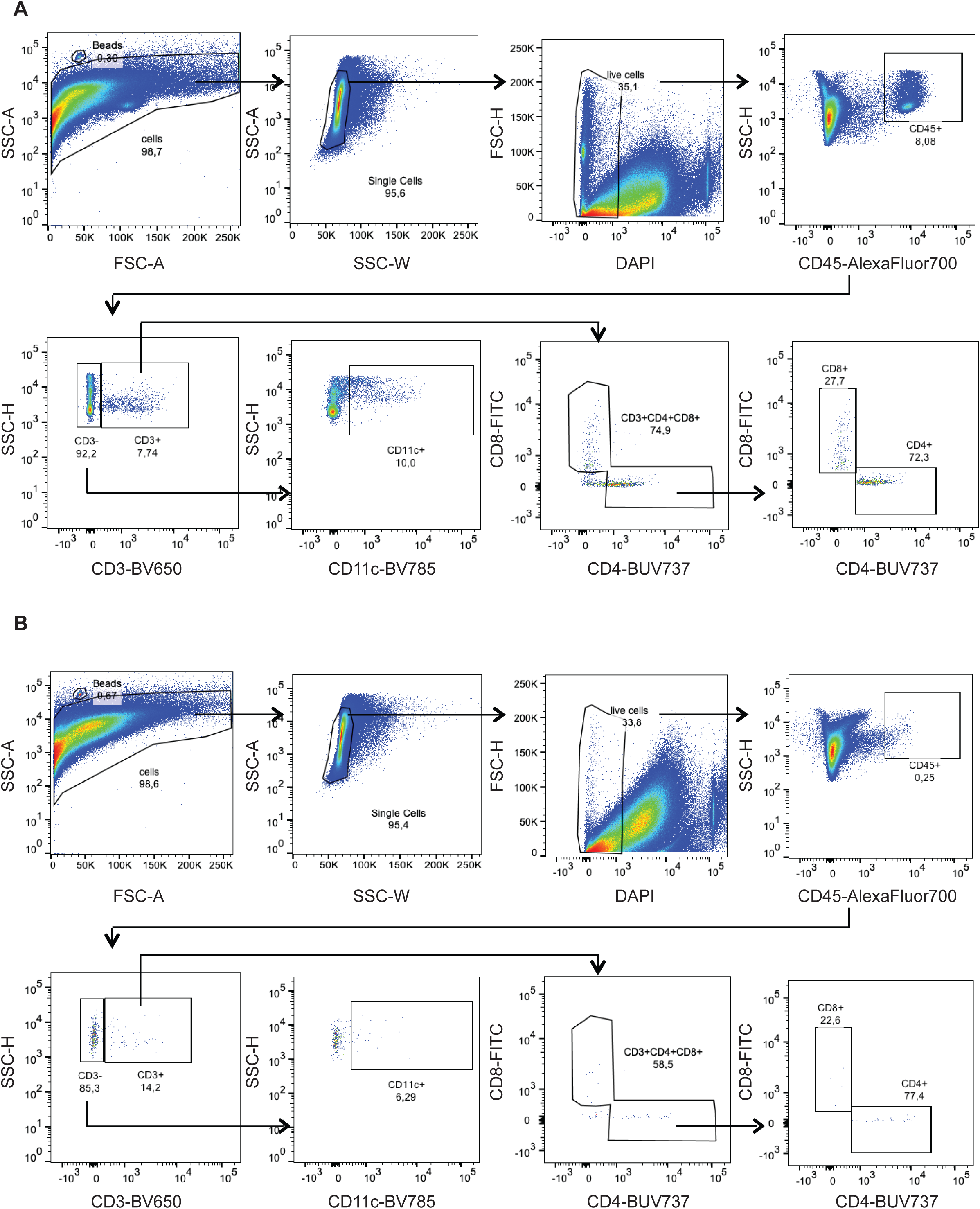
(A-B) Representative flow cytometric analysis of MSI gastric tumors after treatment with antibodies targeting CTLA-4 (A) or IgG control (B). Placement of gates was based on FMO controls.

